# Characterization of 3D organotypic epithelial tissues reveals tonsil-specific differences in tonic interferon signaling

**DOI:** 10.1101/2023.01.19.524743

**Authors:** Robert Jackson, Esha V Rajadhyaksha, Reid S Loeffler, Caitlyn E Flores, Koenraad Van Doorslaer

**Author notes:** Corresponding author: Koenraad Van Doorslaer.

## Abstract

Three-dimensional (3D) culturing techniques can recapitulate the stratified nature of multicellular epithelial tissues. Organotypic 3D epithelial tissue culture methods have several applications, including the study of tissue development and function, drug discovery and toxicity testing, host-pathogen interactions, and the development of tissue-engineered constructs for use in regenerative medicine. We grew 3D organotypic epithelial tissues from foreskin, cervix, and tonsil-derived primary cells and characterized the transcriptome of these *in vitro* tissue equivalents. Using the same 3D culturing method, all three tissues yielded stratified squamous epithelium, validated histologically using basal and superficial epithelial cell markers. The goal of this study was to use RNA-seq to compare gene expression patterns in these three types of epithelial tissues to gain a better understanding of the molecular mechanisms underlying their function and identify potential therapeutic targets for various diseases. Functional profiling by over-representation and gene set enrichment analysis revealed tissue-specific differences: *i.e.*, cutaneous homeostasis and lipid metabolism in foreskin, extracellular matrix remodeling in cervix, and baseline innate immune differences in tonsil. Specifically, tonsillar epithelia may play an active role in shaping the immune microenvironment of the tonsil balancing inflammation and immune responses in the face of constant exposure to microbial insults. Overall, these data serve as a resource, with gene sets made available for the research community to explore, and as a foundation for understanding the epithelial heterogeneity and how it may impact their *in vitro* use. An online resource is available to investigate these data (https://viz.datascience.arizona.edu/3DEpiEx/).

## INTRODUCTION

Epithelial tissues are found throughout the body and play important roles in various organ systems. The epithelium is multifunctional: providing basic external barrier protection from pathogens (Ohland and MacNaughton, 2010), regulation and secretion of bodily fluids (Frizzell and Hanrahan, 2012), absorption of substances internally and externally (Norris, 1998), and sensation (Slominski et al., 2012). Epithelia can be classified based on the number of cell layers and the shape of the cells (Kurn and Daly, 2022). Tissues involved in absorption, filtration, or diffusion (*e.g.*, the lining of the small intestine and the walls of blood vessels) consist of a single layer of cells (Ohland and MacNaughton, 2010). The more complex stratified epithelia are made up of multiple layers of cells and are found in locations where protection is the primary function. Stratified columnar epithelia have tall, column-shaped cells in the top layer. Stratified cuboidal epithelia have cube-shaped cells in the top layer. They are found in locations that need to withstand moderate mechanical stress, such as the lining of the bronchi in the respiratory system (Tschumperlin et al., 2003). Stratified squamous epithelia have flattened cells in the top layer. They are found in locations that are subjected to wear and tear, such as the skin, the oral cavity, and the female reproductive tract. The skin comprises multiple cell layers, including a layer of stratified squamous epithelia on the outer surface, which helps protect the body from physical trauma, ultraviolet radiation, and infection. The epidermal cells also help regulate body temperature and hydration. The oral mucosa protects the underlying tissues from the abrasive effects of chewing and the acidic environment created by certain foods and drinks (Squier and Kremer, 2001), and in the defense against infection (Slomiany et al., 1996). The cervix is a crucial part of the female reproductive system, serving as the gateway between the uterus and the vagina. The cervix is lined with stratified squamous epithelia that helps protect the underlying tissues from mechanical and chemical stresses. The stratified squamous epithelium of the cervix also produces mucus that helps keep the cervix moist and helps to prevent infections (Gipson et al., 1997).

Although skin, cervix, and oral epithelia are all classified as epithelial tissues, they have distinct embryological origins and developmental pathways. Skin is derived from the ectoderm (Sasai and De Robertis, 1997), the outermost layer of the developing embryo. In contrast, the cervical epithelium is likely derived from the endoderm (Robboy et al., 2017), the innermost layer of the developing embryo. In mice, however, conflicting evidence suggests a potential mesodermal origin (Boutin et al., 1992; Hardy, 2010; Moncada-Madrazo and Rodríguez Valero, 2022). Oral epithelia, including the tonsils, are derived from both the ectoderm and the endoderm (Grevellec and Tucker, 2010; Rossen and Thomsen, 2001).

The stratified epithelia of all these tissues are formed through gradual differentiation from stem-cell-like progenitor cells towards fully differentiated cells at the top of the epithelia (Alonso and Fuchs, 2003). This stratified differentiation is not recapitulated using traditional 2D cell culture approaches. Therefore, differentiation-dependent gene expression cannot be studied using standard cell culture techniques. On the other hand, direct sequencing of biopsies derived from (human) tissues does not specifically isolate the epithelial cells, leading to contamination with other cell types present in the biopsy.

Organotypic 3D epithelial tissue culture methods are techniques used to grow and maintain epithelial cells in a 3D configuration that more closely resembles the *in vivo* tissue architecture and function (Anacker and Moody, 2012; Blanton et al., 1991; Dollard et al., 1992; Flores et al., 1999; Meyers, 1996; Meyers et al., 1992; Ozbun and Patterson, 2014). These *in vitro* methods for epithelial stratification were established in the 1970s and further refined over the following decades (Mackenzie and Fusenig, 1983; James G. Rheinwald and Green, 1975). They involve the use of specialized culture systems and media that support the growth and differentiation of cells in a 3D configuration and the formation of functional tissue-like structures. Specifically, we use a collagen gel matrix to support the growth of cells in 3D (Bell et al., 1983, 1981, 1979). Collagen is a structural protein found in many tissues and organs. It is commonly used in tissue culture because it is biocompatible and can support the growth and differentiation of many cell types (Parenteau-Bareil et al., 2010). Primary, host biopsy-derived epithelial cells are cultured to isolate epithelial cells (J. G. Rheinwald and Green, 1975) and homogeneous populations of cells are grown on top of a collagen gel matrix mixed with human fibroblasts. Cells are cultured at the air-liquid interface to promote differentiation. Organotypic 3D epithelial tissue culture methods have several applications, including the study of tissue development and function (Smith et al., 2022), drug discovery and toxicity testing (Markus et al., 2021), host-pathogen interactions (Chatterjee et al., 2019; Israr et al., 2018; Klymenko et al., 2017; Ma et al., 2022; Mole et al., 2009; Roberts et al., 2019), and the development of tissue-engineered constructs for use in regenerative medicine (Schweiger and Jensen, 2016).

We chose a reductionist approach to carefully characterize and contrast the tissue-specific transcriptome of primary epithelial cells (foreskin, cervix, and tonsils) differentiated into stratified squamous epithelia. The use of 3D organotypic rafts in conjunction with RNA-seq represents a significant innovation in the study of epithelial tissues, as it enables a more comprehensive and accurate assessment of gene expression patterns (Hoang et al., 2014; Marioni et al., 2008). By characterizing multiple independent donors of distinct-origin epithelial cells, differentiated using an identical 3D raft culture method (Jackson et al., 2020b), we focus on these tissues’ intrinsic biological similarities and differences. Using this approach, we compare gene expression patterns in these three types of epithelial tissues to better understand the molecular mechanisms underlying their function and identify potential therapeutic targets for various diseases. By characterizing these tissues, we provide a valuable resource for the research community in utilizing origin-specific epithelial tissue for *in vitro* experiments.

This study compared epithelial cells derived from foreskin, cervical, and tonsillar tissues. Despite their different origins, the epithelial cells within these tissues are involved in responding to infection (Nestle et al., 2009). PAMPs, or pathogen-associated molecular patterns, are molecules associated with pathogens (such as viruses, bacteria, fungi, and parasites) and are recognized by pattern recognition receptors (PRRs) expressed by host cells. The recognition of PAMPs by PRRs triggers the activation of the immune response, including the production of type I interferons. Interferons are produced in response to viral infections or other types of cellular stress. Interferons play a vital role in the immune response by activating antiviral gene expression and inhibiting virus replication (Schoggins, 2019). Specifically, interferon binds to receptors on the surface of neighboring cells, activating a signaling pathway culminating in the expression of interferon-stimulated genes.

Interestingly, a subset of these interferon-stimulated genes is continuously expressed without an acute interferon signal (Mostafavi et al., 2016). It is thought that these tonic interferon responses play a role in maintaining immune surveillance and protecting against viral infections. The tonsils play a vital role in the immune system, serving as the first line of defense against infections by trapping and neutralizing pathogens that enter the body through the oral cavity (Siegel, 1984). Tonsillar epithelia are equipped with a variety of immune defense mechanisms. Some evidence suggests that the microenvironment in the tonsils may be different from that of other tissues (Michea et al., 2013; Vangeti et al., 2019; Wu et al., 2011). Tonsils and other lymphoid organs, such as the spleen and lymph nodes, are characterized by a high density of immune cells (Lettau et al., 2020) and a complex network of blood vessels (Ager, 2017). By analyzing the extensive transcriptome data provided as part of this resource, we suggest that the epithelial cells play a critical role in shaping the immune microenvironment to balance tissue homeostasis and immune responses in the tonsils.

## METHODS

### Primary cell culture

Primary human keratinocytes used in these experiments were derived from single donor *ex vivo* explants of neonatal foreskin (Lace et al., 2014), hysterectomy-derived ectocervix (Sprague et al., 2002), and tonsillectomy-derived palatine tonsils (Jackson et al., 2020b). Foreskin and tonsil samples were provided by Banner University Medical Center Tucson as approved by The University of Arizona Institutional Review Board. Primary cervical cultures were kindly provided by Dr. Aloysius Klingelhutz (Sprague et al., 2002). We used three independent donor cultures for each tissue origin for these experiments. Isolated primary keratinocytes were maintained as monolayer co-cultures with irradiated mouse J2 fibroblast feeder cells (J. G. Rheinwald and Green, 1975). To preserve their lifespan, primary epithelial cells were conditionally and reversibly immortalized using the Rho-kinase inhibitor Y-27632 at a 10 μM concentration (Chapman et al., 2014, 2010). Y-27632 was removed prior to growing the cells as 3D organotypic raft cultures. Y-27632 has no impact on the ability of primary cells to differentiate using the raft method (Chapman et al., 2010).

### 3D organotypic raft culture

Full-thickness 3D organotypic epithelial raft cultures were grown over two weeks to yield stratified squamous epithelium (Jackson et al., 2020b; Ozbun and Patterson, 2014). A pair of raft cultures, one for RNA and one for histology, were grown for each of the nine independent donor cultures: *n* = 3 for each tissue origin; foreskin, cervix, and tonsil. Rafts were constructed by first creating a dermal equivalent, where hTERT-immortalized neonatal human foreskin fibroblasts (HFF-hTERT cells) were embedded (8 x 10^4^ per raft) into a solidified rat tail type I collagen matrix. Three consecutive batches of dermal equivalents were made. On the next day, primary human epithelial cells were seeded at equal proportions (2.5 x 10^5^ per raft) atop the dermal equivalents, with representation from each of the tissue origins across each dermal batch, to create confluent basal layers. Finally, on the third day, cultures were lifted onto a hydrophilic polytetrafluoroethylene membrane insert (0.4 μm pore size) for an air-liquid interface and fed every two days (seven total media changes) with differentiation media (1.88 mM Ca^2+^ and no epidermal growth factor, EGF) to stratify until fully differentiated raft cultures were harvested on day 15 post-lift.

### Histology

Whole organotypic epithelial raft cultures were fixed in 10% neutral-buffered formalin for 24 hrs at room temperature then gently washed and stored in 70% ethanol. Fixed rafts were bisected, paraffin-embedded, sectioned (4 µm thickness), and hematoxylin and eosin (H&E) stained at the University of Arizona Cancer Center’s Acquisition and Cellular/Molecular Analysis Shared Resource (TACMASR). Color micrographs were captured using an Olympus BX50 microscope, DP72 camera, and cellSens Entry software. Whole palatine tonsils were collected from Banner University Medical Center Tucson as described above and histologically processed in the same manner as organotypic raft cultures. Overlapping edge color micrographs were captured using an EVOS M5000 microscope and stitched together using Microsoft Image Composite Editor.

### Immunofluorescence

Protein-level characterization was performed as described (Jackson et al., 2020b). Unstained slides with formalin-fixed paraffin-embedded epithelial sections (4 µm thick) were deparaffinized by xylenes and ethanol washes. Slides were steamed for 30 min in 10 mM citrate buffer (pH 6), blocked with 2% bovine serum albumin (BSA) in phosphate-buffered saline (PBS) for 30 min, and incubated overnight at 4°C with primary antibodies diluted in blocking buffer. The following primary antibodies and conditions (dilutions and final concentrations) were used: anti-COL17A1 basal epithelial marker (rabbit polyclonal, 1:500, 0.6 µg/mL, MilliporeSigma, Catalog No. HPA043673-100UL) and anti-CRNN upper epithelial marker (goat polyclonal, 1:200, 1 µg/mL, Novus Biologicals, Catalog No. 103242-512). Following PBS washes, slides were incubated in the dark for 30 min at room temperature with secondary antibodies diluted 1:800 in blocking buffer: donkey anti-rabbit AlexaFluor 488 (Thermo Fisher Scientific, Catalog No. A-21206) and donkey anti-goat AlexaFluor 555 (Thermo Fisher Scientific, Catalog No. A-21432). Slides were PBS washed, DAPI-stained (1:1000), ddH_2_O rinsed, mounted with ProLong Diamond Antifade Mountant (Thermo Fisher Scientific, Catalog No. P36965), and imaged by epifluorescence using a Leica DMIL LED microscope, DFC3000G camera, and Leica Application Suite X software. For each set of images taken, the DAPI (blue), COL17A1 (green), and CRNN (red) channels were merged in Adobe Photoshop. Reference images were obtained from The Human Protein Atlas (v21.proteinatlas.org) for skin, cervix, and oral mucosa. Quantification of midzone gap distance, which represents the double-negative space between CRNN and COL17A1 layers, was performed in ImageJ/Fiji by measuring perpendicularly from the inferior portion of the CRNN-positive area to the superior portion of the basal layer. Measurements were taken at 25 μm intervals spanning the length of the tissue within each field of view. To determine relative differences due to tissue origin we calculated midzone gap ratios by dividing each midzone gap thickness measurement by the mean of the donor sample of the smallest gap within each set of tissues (T3 for 3D raft cultures and oral mucosa 1505 for the reference tissues).

### RNA-seq

RNA was isolated from the epidermal compartment (containing only basal and suprabasal keratinocytes) of organotypic raft cultures using Qiagen’s RNeasy Mini Kit (Catalog No. 74106) after separating them with sterile forceps from their underlying dermal equivalents and flash-freezing with liquid nitrogen (Jackson et al., 2020b). RNA was eluted in nuclease-free water, carry-over DNA was removed with the TURBO DNA-free Kit (Thermo Fisher Scientific, Catalog No. AM1907), then shipped on dry ice to Novogene Corporation (Sacramento, CA). High-quality eukaryotic mRNA libraries were prepared and sequenced on an Illumina platform, yielding >20 million paired-end reads (2x150 bp length) per sample. Raw sequence reads were deposited in the Sequence Read Archive (SRA) as BioProject ID PRJNA916745. High-throughput sequencing data were initially processed using the CyVerse Discovery Environment (http://www.cyverse.org), where FastQC (https://www.bioinformatics.babraham.ac.uk/projects/fastqc/) was used for quality screening, HISAT2 (Kim et al., 2015) for mapping sequences to the human reference genome version GRCh38, and featureCounts (Liao et al., 2014) for generating gene-level read counts. Principal component and differential expression analysis of read counts, including library-size correction and statistical testing, was performed using the DESeq2 Bioconductor package (Love et al., 2014) implemented in R (https://www.R-project.org/) via RStudio’s integrated development environment (http://www.rstudio.com/). PcaExplorer was used to plot 95% confidence interval ellipses for the principal components (Marini and Binder, 2019). The 18,925 confidently detected genes were identified using default DESeq2 parameters and were a subset of 31,033 genes with non-zero total read counts among the 58,884 total Ensembl gene IDs used for mapping. Lists of differentially expressed genes, with a stringent adjusted *P*-value < 0.01, were analyzed for functional enrichment using g:Profiler (Kolberg et al., 2020; Raudvere et al., 2019). Unsupervised clustered heatmaps were constructed in R using the pheatmap package (https://cran.r-project.org/package=pheatmap). Pairwise contrast volcano plots were made using EnhancedVolcano (https://github.com/kevinblighe/EnhancedVolcano). Gene set enrichment analyses (GSEA) and over-representation analysis of pairwise and overlapping genes were done using clusterProfiler (Wu et al., 2021; Yu et al., 2012). Area-proportional ellipse Euler diagrams were made using eulerr (https://CRAN.R-project.org/package=eulerr). The singscore package (Bhuva et al., 2020; Foroutan et al., 2018) was used to calculate tissue-identifying gene expression signature scores. The web-based submission tool for Integrated Motif Activity Response Analysis (ISMARA) was used to identify regulatory motifs and transcription factors that regulate gene expression (Balwierz et al., 2014).

### Reanalysis of Mostafavi et al., dataset

Mostafavi and colleagues used Affymetrix ST1.0 microarrays to measure the expression of interferon-stimulated genes in mouse B-cells and macrophages (Mostafavi et al., 2016). They report the expression of each gene at baseline and 2 hours post interferon stimulation in wildtype (WT) mice. To identify genes that do not require interferon, the experiments were repeated in interferon receptor (IFNAR1) KO mice (KO). Supplemental data S1 provided by Mostafavi and colleagues was downloaded and further analyzed. If a microarray probe was detected in both macrophages and B-cells, the expression values were averaged. Otherwise, the B-cell or macrophage value was used. For some genes, multiple probes were present, for simplicity, only one probe per gene was analyzed. For some genes, the human homologue could not be identified. Similarly, some of the mouse genes were auto-converted to “dates” by Microsoft Excel and could not be recovered (Ziemann et al., 2016). These genes were excluded from our further analyses.

For each gene, a fold change of baseline vs. IFNAR1-KO (WT/KO) expression and the intrinsic responsiveness to interferon (WT+IFN / WT) were calculated. As per Mostafavi and colleagues, highly tonic-sensitive genes are those genes where log_2_(foldchange(WT/KO))>log_2_(foldchange(WT+IFN / WT))/4. With low tonic-sensitive genes following log_2_(foldchange(WT/KO))<log_2_(foldchange(WT+IFN / WT))/6. We consider the genes that are larger than log_2_(foldchange(WT+IFN / WT))/6 but lower than log_2_(foldchange(WT+IFN / WT))/4 to have medium sensitivity. The data and data manipulation steps are available in the supplementary materials associated with **Figure 9**.

### 3DEpiEx application development

The three-dimensional (<u>3D</u>) <u>Epi</u>thelial Gene <u>Ex</u>pression (3DEpiEx) application was developed using the Shiny R package (Kasprzak et al., 2020): https://shiny.rstudio.com/. Application development included two main parts: the user interface and the server. The user interface dictates user interaction and the overall appearance of the application, while the server involves handling user inputs and the construction of different outputs. The primary input for 3DEpiEx is the user gene selection. With nearly 19,000 genes from which to choose, it was of the utmost importance to develop an effective tool that would allow users to select specific genes based on Ensembl ID (Cunningham et al., 2022) or HGNC gene symbol (Tweedie et al., 2021). Ultimately, the “selectizeInput” search method (https://shiny.rstudio.com/articles/selectize.html), which combines a search bar and a dropdown menu, was utilized. The biomaRt library (Smedley et al., 2015) was used to annotate gene symbols for the pre-existing data which already contained Ensembl IDs. Once the gene selection tool was created, outputs for the box plots and volcano plots were implemented via volcano3D: https://cran.r-project.org/web/packages/volcano3D/vignettes/Vignette.html. The last part of the development process was web hosting and publication via RStudio Connect. This allowed for public access and use of the application, freely available at https://viz.datascience.arizona.edu/3DEpiEx/.

## RESULTS

Three-dimensional organotypic raft cultures are the current state-of-the-art method to reconstruct fully stratified 3D epithelial tissue *in vitro* (Andrei et al., 2010; Arnette et al., 2016; Jackson et al., 2020b; Roth-Carter et al., 2022). Traditionally, most of these 3D tissues are grown from epithelial cells derived from foreskin keratinocytes (Andrei et al., 2010; Arnette et al., 2016; Frattini et al., 1996; Kesimer et al., 2010). However, since epithelial tissues have different embryogenic origins, these foreskin-derived tissues may not be representative of other (squamous) epithelia. Therefore, we designed this comparative study to characterize 3D tissues derived from primary cells obtained from foreskin, cervical, or tonsillar biopsies.

### *In vitro* 3D organotypic epithelia derived from isolated primary cells recapitulate *in vivo* histology

Primary epithelial cells obtained from foreskin, cervical, or tonsil tissue were cultured to generate 3D organotypic rafts as described (Jackson et al., 2020b). For each tissue, cells from three unique donors were included. These nine 3D organotypic epithelial raft cultures were grown using the identical method and materials as described (Jackson et al., 2020b). All nine 3D organotypic raft cultures were cultured at the same time using the identical reagents. Rafts were processed in sets of three samples, one sample for each tissue. Therefore, any observed differences represent inherent epithelial cell properties (*e.g.*, tissue origin and donor genetics) and are not due to differences in culture methods. Following stratification, we sectioned and stained formalin-fixed tissues with hematoxylin and eosin (H&E; **Figure 1A**). All cultures (*i.e.*, 3 donors and 3 tissue origins) differentiated into fully stratified 3D organotypic epithelia **(Figure 1–figure supplement 1**). We identified basal keratinocytes at the dermal interface (dotted white line in **Figure 1A**), suprabasal midzone cells, and differentiated squamous keratinocytes at the apical surface of the tissue. Despite having used identical culturing methods, we observed tissue-specific differences between samples. Compared to foreskin-derived tissues, the suprabasal midzone cells of the mucosal organotypic epithelial tissue (cervix and tonsil) had a greater extent of cytoplasmic clearing. This cytoplasmic clearing indicates high glycogen content (Falin, 1961). Cytoplasmic glycogen (and other neutral mucins) are rapidly degraded post-harvest and during the formalin fixation process (Sullivan et al., 2015) and poorly stained by H&E (Meyerholz et al., 2018). Indeed, abundant cytoplasmic glycogen is typical of the normal ectocervical epithelium (Gregoire et al., 1973).

**Figure 1.**
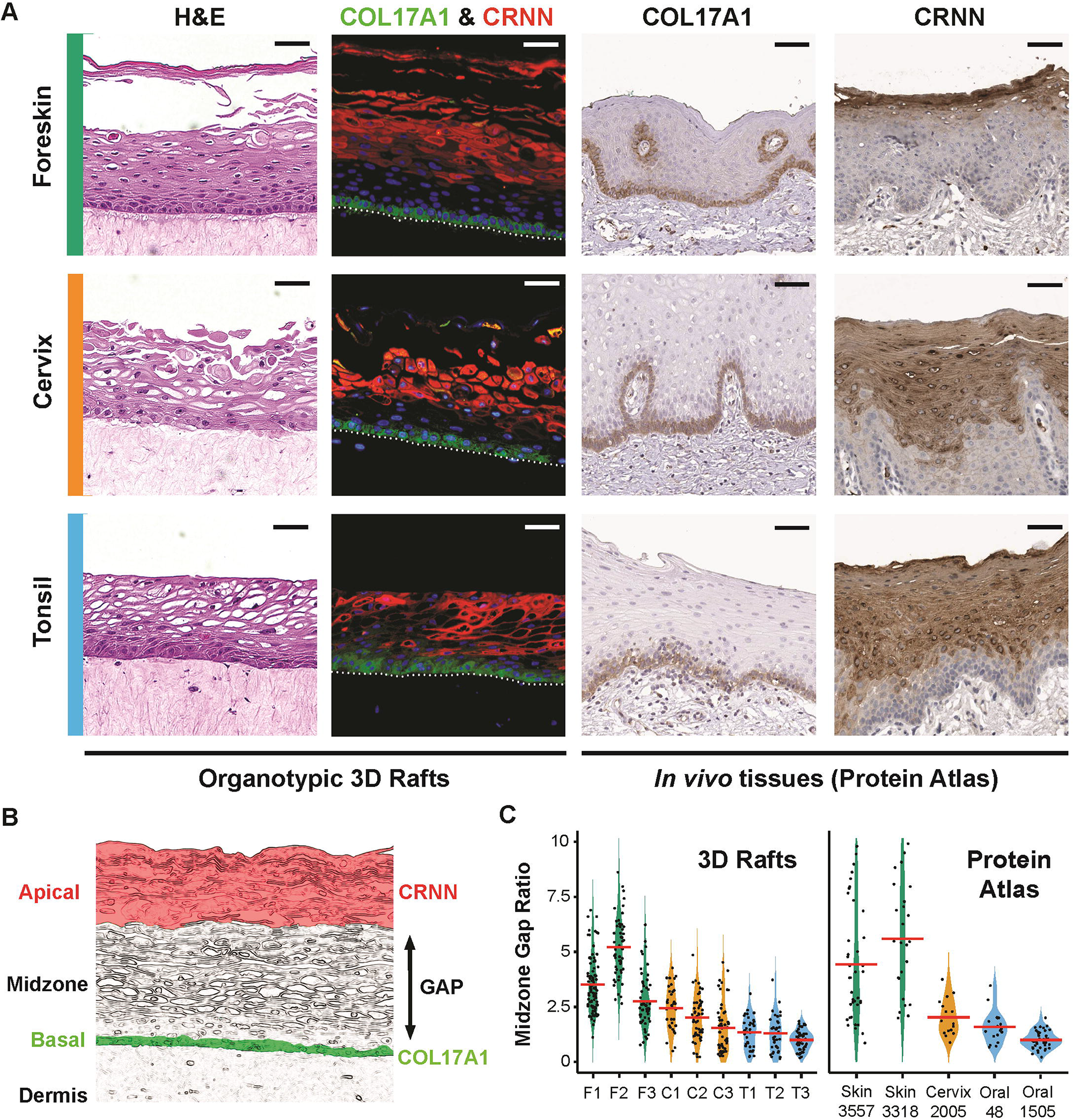
Histological and immunofluorescent verification of multi-origin 3D organotypic raft cultures. **(A)** The top row represents foreskin epithelia, the middle row represents cervical epithelia, and the bottom row represents oral epithelia. The first column shows H&E micrographs of 3D organotypic epithelial raft cultures. The second column depicts immunofluorescent staining of the 3D raft cultures with COL17A1 (Collagen XVII; green), CRNN (Cornulin; red), and DNA (DAPI-stained; blue). Columns three and four contain immunohistochemical reference staining of COL17A1 and CRNN from The Human Protein Atlas. Colorimetric 3,3’-diaminobenzidine (DAB)-stained COL17A1 micrographs include: normal skin tissue (T-X0500, Female, age 66, Patient id: 4786), normal cervical tissue (T-83000, Female, age 37, Patient id: 4504), and normal oral mucosa tissue (T-51000, Male, age 62, Patient id: 3724) all stained with antibody HPA043673. CRNN micrographs include: normal skin tissue (T-80100, Female, age 66, Patient id: 3357), normal cervical tissue (T-83000, Female, age 55, Patient id: 2005), and normal oral mucosa tissue (T-51000, Male, age 62, Patient id: 1505) all stained with antibody CAB026182. Images are available from v21.proteinatlas.org. Scale bars represent 50 μm. **(B)** Schematic illustration of the apical, midzone, basal, and dermal layers in stratified squamous epithelia. CRNN-positive cells are typically present in the apical layer (highlighted in red), while COL17A1-positive cells are typically present in the basal layer (highlighted in green). The midzone gap represents the double-negative area from the inferior portion of CRNN to the superior portion of basal epithelial cells. **(C)** Midzone gap distances were quantified between skin, cervical, and oral epithelia in our 3D rafts and reference Protein Atlas images (patient id indicated for each tissue, v21.proteinatlas.org). Violin plots were made for each independent donor measured while points represent the ratio of each midzone gap distance measurement divided by the lowest donor’s average midzone gap measurement. All measurements were taken at 25 μm intervals spanning the length of visible tissue across non-overlapping fields of view using ImageJ/Fiji. Mean ratios for each donor are indicated by red bars.

We used immunofluorescence to characterize these tissues further. We stained these tissues with antibodies specific for collagen XVII A1 (COL17A1) and cornulin (CRNN). COL17A1 is expressed in epithelial hemidesmosomes, is a known basal cell marker in stratified squamous epithelial tissue, and is an integral component of the basement membrane (Christiano and Uitto, 1996). COL17A1 protein was detected in cells localized to the basal layer that contact the basal membrane at the dermal interface (**Figure 1A; green**). Suprabasal cornulin is a terminal differentiating cell marker in the superficial layers of differentiated epithelial tissue (Bedard et al., 2021; Contzler et al., 2005). We detected cornulin in the superficial layers of 3D organotypic epithelial raft cultures (**Figure 1A; red**). All tissues have a section of double-negative midzone cells (COL17A1^NEG^/CRNN^NEG^) representing a clear distinction between the basal COL17A1(+) monolayer and apical CRNN^POS^ layers of these tissues (**Figure 1B**). Comparing the height of the midzone gap between the tissues (**Figure 1C**), the gap was most prominent in foreskin-derived 3D epithelia (32 +/-11 μm, *n* = 3 tissues), moderate in cervix-derived cultures (17 +/-4 μm, *n* = 3 tissues), and smallest in tonsil-derived cultures (10 +/-2 μm, *n* = 3 tissues). We identified a similar pattern of COL17A1^NEG^/CRNN^NEG^ midzone gap heights (**Figure 1C**) in reference micrographs obtained from The Human Protein Atlas (v21.proteinatlas.org) tissue database (Uhlén et al., 2015). The midzone gap measured between the COL17A1^POS^ basal cells and the CRNN^POS^ apical cells was most prominent in the skin (69 +/-10 μm, *n* = 2), moderate in the cervix (29 μm, *n* = 1), and smallest in the oral mucosa (18 +/-6 μm, *n* = 2). Since 3D organotypic rafts have a reduced total epithelial thickness compared to *in vivo* tissue, we compare the midzone gap thickness to the oral raft or tissue. The midzone gap in skin-derived tissues is roughly 3 times as wide as those in oral-derived tissue (3.1x 3D rafts and 3.9x for skin). For both cervical rafts and cervical tissue the midzone gap is 1.6x wider than the oral equivalent (**Figure 1C**).

Tonsillar epithelia are composed of reticular crypt epithelia and surface epithelia (**Figure 2**) (Clark et al., 2000). The reticular crypt epithelium likely plays an important role in regulating the immunological functions of the tonsil (Perry, 1994). The crypt epithelium also has reduced barrier function, while the surface epithelia, like the surrounding oral mucosa, is thought to serve a typical epithelial barrier role owing to tight junctions (Go et al., 2004). When cultured as 3D organotypic rafts, two of the tissues (T1 and T2) display reticular/crypt epithelial characteristics, including a reduced overall thickness and extent of stratification and a spongey-like appearance (**Figure 2**). The T3 sample more closely resembles surface epithelium, with more layers of stratified squamous cells and signs of apical keratinization/cornification. Compare the micrograph derived from a tonsillectomy to the H&E-stained 3D organotypic rafts (**Figure 2**): focusing on the thickness differences between crypt and surface epithelia, as well as the extent of stratification. Also, note the extent of lymphocyte infiltration in the *ex vivo* crypt epithelia and the reticulated “spongey-like” appearance.

**Figure 2.**
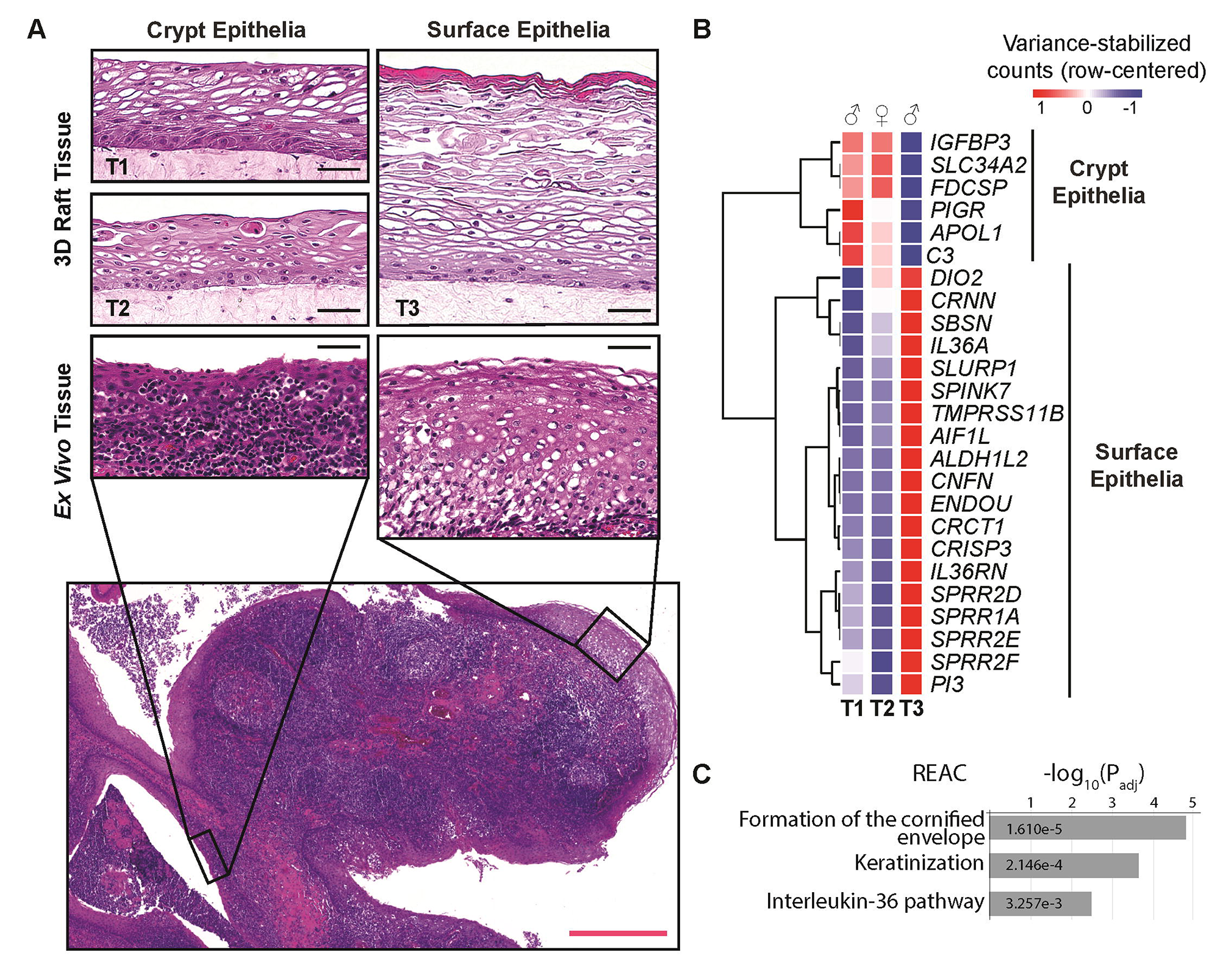
3D organotypic raft cultures recapitulate differences between crypt- and surface-like tonsillar epithelia. **(A)** Histology of crypt-like and surface-like tonsillar epithelia. The top row contains H&E micrographs of three donor-derived 3D tonsillar epithelial raft cultures (T1, T2, and T3). Below we show micrographs from real tissue: a stitched composite of the human palatine tonsil, which includes different types of tonsillar epithelia: crypt epithelia on the left and surface epithelia on the right. The surface epithelium of tonsils consists primarily of stratified squamous cells and make up the outer periphery, whereas crypt epithelium serves to increase the overall surface area and is composed of an uneven mixture of stratified squamous cells and non-epithelial cells (*e.g.*, infiltrating lymphocytes), resulting in a spongy appearance. As a result of these epithelial subtypes, there is a variable expression of differentiation-related epithelial markers in tonsillar tissues. Black scale bars represent 50 μm and the red scale bar represents 500 μm. **(B)** Transcriptional variation between surface-like (T3) and crypt-like (T1, T2) 3D tonsillar epithelia: top 25-varying genes within tonsil samples. **(C)** Over-representation analysis of surface-like tonsillar epithelia markers (REAC = Reactome pathway).

In conclusion, 3D organotypic raft cultures recapitulate several histological features of the *in vivo* tissues from which the cells were initially derived.

### Epithelial origin is the primary determinant of transcriptomic variability between foreskin, cervix, and tonsil-derived 3D rafts

We analyzed the transcriptome of 3D organotypic epithelial tissues using RNA-seq (**Supplementary File 1, 2**) to compare and contrast foreskin, cervix, and tonsil-derived tissues. We carefully peeled the epidermis off the dermal equivalent and isolated total RNA from these cells. Poly-A enriched cDNA was sequenced by a commercial provider using an Illumina NovaSeq. **Table 1** shows that sequencing depth (average ∼29 million paired-end reads per sample) and quality (∼98% of sequences had at least a Q30 quality score) were sufficient for further analyses (Ewing and Green, 1998; Liu et al., 2014). Roughly 97% of the paired-end sequencing reads for each sample were mapped to the human reference (GRCh38, **Table 2**) using HISAT2 (Kim et al., 2015). The library-size correction was performed using DESeq2 (Love et al., 2014) (**Supplementary File 3**). Cook’s distances for all samples were inspected via violin plots to ensure no individual samples were overtly contributing to outliers (**Figure 3–figure supplement 1A**). From the 31,033 genes with detectable counts, 12,038 were excluded due to low counts (mean count < 3), and 70 were excluded as count outliers (based on Cook’s distance, **Figure 3–figure supplement 1B**).

**Table 1.** RNA-seq file quality summary. Filename, Size (in bytes), and MD5 checksum were used for verifying file identity and integrity. FastQC was run on each each sequencing read file to verify quality. The percentage of sequences with quality scores of at least 30 (≥Q30%, 1 in 1000 probability of an incorrect base call, 99.9% inferred base call accuracy) was calculated from the raw “fastqc_data.txt” output for each file.

| Filename | Size (Bytes) | MD5 Checksum | Total Sequences | $\geq Q30\%$ |
| --- | --- | --- | --- | --- |
| F1_1.fq.gz | 2,088,551,236 | 8aa04a99fecc398bdd8a9a50dd9eb7ed | 28,278,312 | 97.85 |
| F1_2.fq.gz | 2,129,194,266 | 3f3ff98b84bf43be00efe178a34a45ed | 28,278,312 | 97.83 |
| F2_1.fq.gz | 2,272,389,773 | 492556bef87269c16bf316cafb721126 | 30,553,465 | 97.63 |
| F2_2.fq.gz | 2,317,151,216 | 38060cac3de2d83672922be17e0c1f0e | 30,553,465 | 97.59 |
| F3_1.fq.gz | 1,845,676,286 | 62be31900625b5e817459c9922228774 | 24,879,571 | 97.72 |
| F3_2.fq.gz | 1,880,871,095 | 11c0c35f4ef31c16bb67707db474c89b | 24,879,571 | 97.72 |
| C1_1.fq.gz | 2,474,035,817 | 828128f19d2c1f963c6ea1a78c1b303b | 33,324,902 | 97.94 |
| C1_2.fq.gz | 2,505,869,077 | 2d46b65df75822a91814fedf0114ca00 | 33,324,902 | 98.13 |
| C2_1.fq.gz | 2,282,090,096 | 22687d29ebe6d3d883fba4708acdb6d5 | 30,857,610 | 97.70 |
| C2_2.fq.gz | 2,319,433,070 | 467c52ca7173bc734089abad20077ea3 | 30,857,610 | 97.74 |
| C3_1.fq.gz | 1,691,719,201 | fad9ded3da55dab63f958d9c3baa9968 | 22,735,206 | 97.64 |
| C3_2.fq.gz | 1,726,093,353 | 3f7187eefc7d50927d99da10c880e0ee | 22,735,206 | 97.58 |
| T1_1.fq.gz | 2,435,070,557 | 2e2132cd1c50a04b50228d8693edb16f | 32,788,230 | 97.85 |
| T1_2.fq.gz | 2,477,227,739 | 0b5618e1499e87f44b7c384e0e95fb38 | 32,788,230 | 97.87 |
| T2_1.fq.gz | 2,495,864,252 | f7725f7d69d133932bf8b5dc398430b1 | 33,805,463 | 97.86 |
| T2_2.fq.gz | 2,551,617,978 | cd2ca3c8b7bd23577725b9d8a5c52a0b | 33,805,463 | 97.82 |
| T3_1.fq.gz | 1,583,107,240 | 1cbfb4be79d2431a4e09c8d46f26b197 | 21,433,198 | 97.83 |
| T3_2.fq.gz | 1,610,013,928 | 464026c424aa564cdb223bfd48ea3c27 | 21,433,198 | 97.86 |

**Table 2.** RNA-seq mapping and gene-level assignment summary. Homo sapiens (GRCh38) mapped reads comprised 96.72 +/-0.20% of the total reads via HISAT2 (mapped + unmapped). FeatureCounts was used for gene-level counting of the HISAT2 alignment output through a series of category filters (left to right, starting with human gene “assigned”, which is used for downstream differential expression analyses).

| **Sample** | **Human %** | **Assigned %** | **Unmapped %** | **Multi Mapping %** | **No Features %** | **Ambiguity %** |
| --- | --- | --- | --- | --- | --- | --- |
| F1 | 96.84 | 68.93 | 1.81 | 21.50 | 2.84 | 4.92 |
| F2 | 96.77 | 68.54 | 1.78 | 21.16 | 3.42 | 5.10 |
| F3 | 96.93 | 69.94 | 1.68 | 20.04 | 3.10 | 5.24 |
| C1 | 96.92 | 72.66 | 1.95 | 17.39 | 2.55 | 5.45 |
| C2 | 96.75 | 69.02 | 1.92 | 20.44 | 3.24 | 5.38 |
| C3 | 96.34 | 69.83 | 2.27 | 17.44 | 5.33 | 5.14 |
| T1 | 96.74 | 71.81 | 1.97 | 18.21 | 2.69 | 5.32 |
| T2 | 96.79 | 67.30 | 1.94 | 22.59 | 3.26 | 4.92 |
| T3 | 96.44 | 69.72 | 2.19 | 20.64 | 2.62 | 4.84 |

As an initial analysis, we used principal components to analyze the top 500 varying genes (based on gene-level counts). Principal component analysis revealed that each tissue had a distinct transcriptional profile (**Figure 3**). The scree plot in **Figure 3A** shows that Principal Component 1 (PC1; 41%) and PC2 (28%) explain 69% of the variance. PC1 and PC2 visually distinguish foreskin (green), cervix (orange), and tonsil (blue) derived tissues from each other (**Figure 3B**), further supporting that 3D organotypic rafts maintain tissue-specific features. The density plots for each tissue along both PC1 and PC2 demonstrate the spatial separation of these transcriptomes along the axes of the individual principal components. We used pcaExplorer (Marini and Binder, 2019) to draw 95% confidence interval ellipses around each set of samples belonging to a specific tissue (**Figure 3B**). The lack of overlap between the ellipses statistically supports that foreskin, cervix, and tonsil-derived tissues have different transcriptional profiles. This separation is further confirmed using a multivariate analysis of variance (MANOVA; *P*-value = 2.89 x 10^-5^).

**Figure 3.**
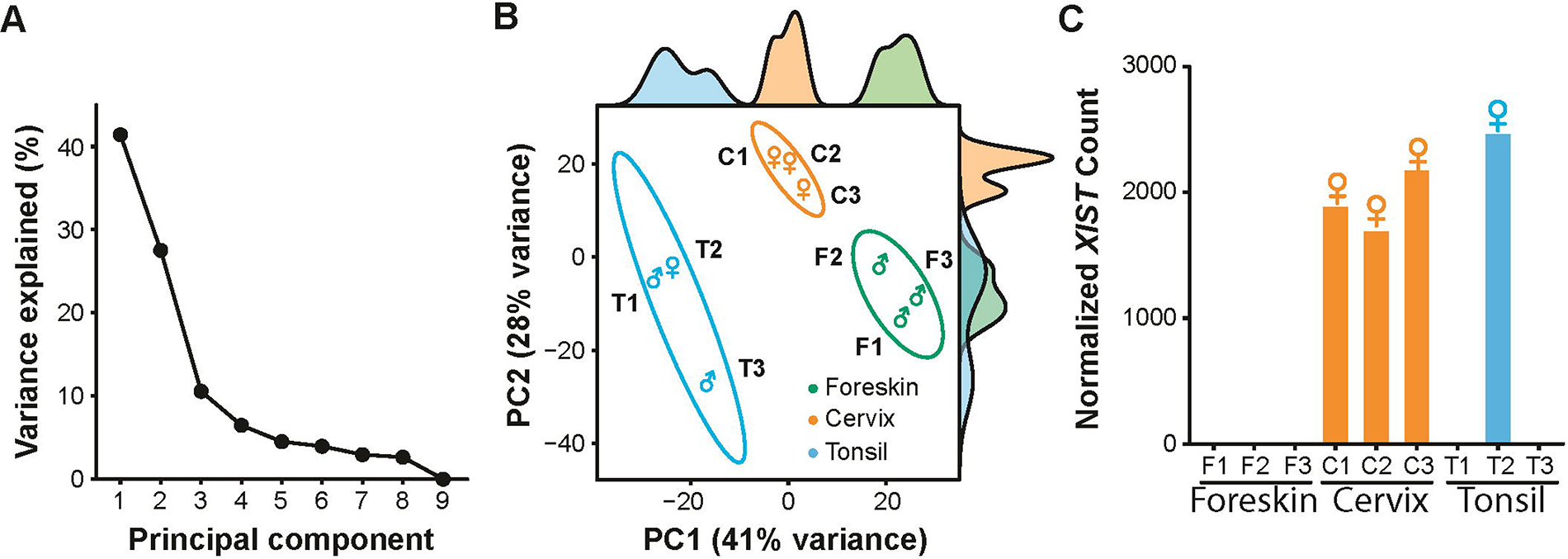
Transcriptomic characterization of multi-origin 3D organotypic raft cultures. **(A)** Scree plot of principal components (PCs) and their variance explained (%), totaling 100%, using the top 500 varying genes (variance-stabilized counts) detected by RNA-seq. PC1 (41%) and PC2 (28%) explained the majority (∼70%) of transcriptional variability. **(B)** Principal component analysis (PCA) plot of 3D organotypic raft culture transcriptomes (top 500 varying genes, variance-stabilized counts), each point represents an individual raft culture from an independent donor, biological sex labeled (♂, male; ♀, female), with 95% confidence ellipses for the tissue origin groups. Density plots for PC1 and PC2 are plotted opposite the respective axis. **(C)** Verification of biological sex by sex-specific marker *XIST*. DESeq2 normalized (library-size corrected) counts for each sample confirmed sample identity.

Importantly, these analyses demonstrate that the tissue origin of keratinocytes, rather than the genetics of the individual donor, is the primary driver of transcriptomic variability of 3D organotypic raft cultures. Of note, biological sex, as determined by the sex-specific expression marker XIST (**Figure 3C**), is not a significant contributor to transcriptional variability among our samples, as T1 (♂, male) clusters closely to T2 (♀, female) (**Figure 3B**).

The densities along PC1 separate each of the tissue groups. Therefore, we conclude that PC1 (41% of the variability) separates the three tissues, whereas PC2 has overlapping distributions and does not clearly distinguish foreskin (green) from tonsil (blue) tissue. This increased spread is likely driven by variability in the tonsillar tissue, *i.e.*, T1/T2 (crypt) vs. T3 (surface). To further investigate this, we plotted the top 25 variable genes between the different tonsil tissues (**Figure 2B**). T3 shows higher expression of genes implicated in tissue keratinization (*e.g.*, CRNN, SBSN, and SPINK7). Indeed, functional profiling of the 19 genes up-regulated in T3 indicates that these genes are implicated in the cornified envelope formation and keratinization (**Figure 2C**). Likewise, T1 and T2 express the follicular dendritic cell-secreted protein (FDCSP) ∼40x higher than T3. FDCSP was previously reported as a marker for crypt epithelia (Marshall et al., 2002; Massoni-Badosa et al., 2022). These data further support the hypothesis that T1/T2 and T3 are derived from different sub-tissues and further confirm that 3D organotypic rafts maintain many features associated with the original, *in vivo* tissue. Nonetheless, as is clear from **Figure 3B**, the tonsillar tissues are transcriptionally more similar to each other than to other tissues. Therefore, we will analyze T1, T2, and T3 as biological replicates.

### Global gene expression across three different tissue-derived 3D rafts

To characterize the tissue-inherent transcriptional differences, we used a likelihood ratio test (LRT) to detect the globally differentially expressed genes between foreskin, cervix, *and* tonsil-derived epithelial. Based on our biological question and supported by the exploratory PCA of the primary sources of variation (**Figure 3B**), we included tissue origin as the grouping factor, with three levels (foreskin, cervix, and tonsil) in the DESeq2 model (Love et al., 2014). The individual donor samples were used as biological replicates (*n* = 3). Using a strict statistical significance threshold (*P*-adj < 0.01), we identified 1,238 globally differentially expressed genes distinguishing the three tissues (**Supplementary File 4**). The combination of different donors and strict statistical cut-off increases the likelihood of identifying biologically relevant differentially expressed genes. These data demonstrate important transcriptional differences between foreskin, cervix, and tonsil-derived tissues. In turn, we may miss important differentially expressed genes due to stringent cut-offs.

We used unsupervised clustering to analyze these differentially expressed genes (**Figure 4**). Unsupervised clustering builds a hierarchy of clusters, which are summarized as a dendrogram. In these heatmaps, individual columns represent the different samples, while differentially expressed genes are shown in rows. The color indicates the relative expression of each differentially expressed gene in the different samples: blue means the gene is expressed less in that sample compared to the normalized gene expression across all samples, whereas red means the gene is overexpressed. Therefore, expression levels should only be compared across samples for the same gene and not between genes. Unsupervised clustering of the samples confirmed that all three tissues are different. Based on this set of genes, the mucosal tissues (cervix and tonsil) are transcriptionally more similar to each other than to foreskin-derived tissues. Hierarchical clustering of the genes identified two main clusters we designate as “HFK-up” and “HFK-down,” respectively (see **Supplementary File 5** for human foreskin keratinocyte, HFK, cluster annotations).

**Figure 4.**
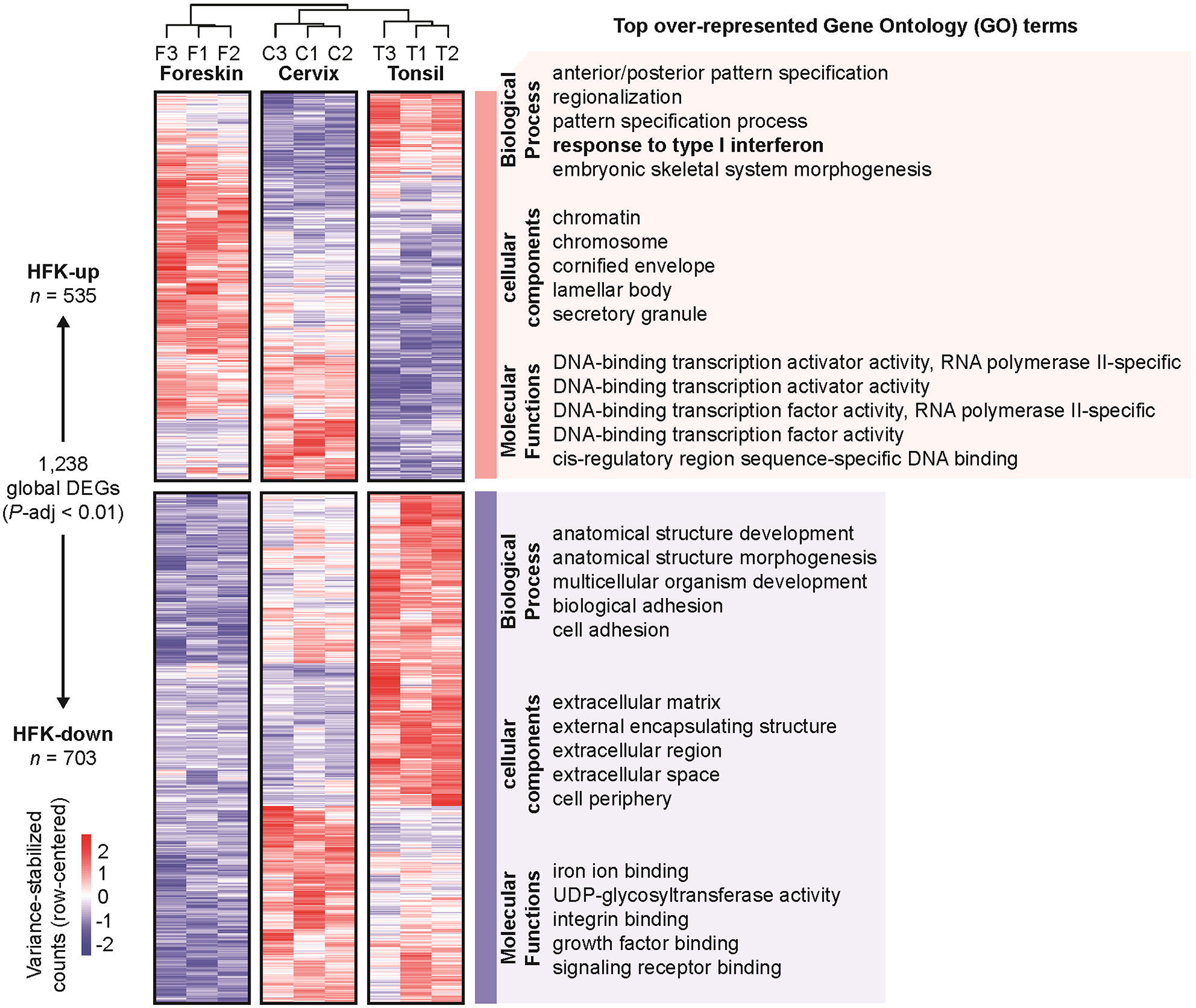
Globally differentially expressed genes. Variance-stabilized, row-centered, counts are shown for all 1,238 genes differentially expressed genes (LRT, *P*-adj < 0.01). For each cluster, the top five over-represented Gene Ontology (GO) terms for each GO category (BP, biological process; MF, molecular function; and CC, cellular component) are listed based on the lowest g:Profiler *P*-adj.

We identified 535 genes up-regulated in foreskin-derived tissues (HFK-up cluster). On the other hand, HFK-down contains 703 genes with reduced steady-state levels compared to the mucosal tissues. Over-representation analysis of the two clusters via g:Profiler (ordered query, g:SCS threshold < 0.05) identified 181 functionally enriched terms for HFK-up (**Supplementary File 6**) and 293 functionally enriched terms for HFK-down (**Supplementary File 7**).

HFK-up genes were enriched for biological processes related to pattern specification and morphogenesis. These embryogenic terms relate to tissue development along an organism’s anterior-to-posterior axis (Frankenberg, 2018). The top enriched cellular components for HFK-up include differentiation-related structures: the cornified envelope and lamellar bodies common to the upper layers of stratified squamous epithelium that are important for barrier maintenance (Menon et al., 1992). The top enriched up-regulated molecular functions in HFKs relate to transcription factor activity. Furthermore, the top enriched cellular components for HFK-up also included chromatin and chromosome subcellular localizations, further supporting the critical role of these DNA-binding transcription factors. While we expected tissue-specific differences in differentiation and transcription factor activity, we were intrigued to find an enrichment of innate immune-related genes and transcription factors supporting the identification of “response to type I interferon” as a top enriched biological process in the HFK-up cluster (**Figure 4**).

HFK-down genes were enriched for differentiation-related biological processes (*e.g.*, development and morphogenesis), further highlighting differences between the tissues related to differentiation. The analyses also identified differences related to cell adhesion processes. Of note, these cell adhesion differences are also reflected by the observations that the top enriched cellular components included the extracellular matrix and cell periphery. Top molecular functions enriched in the HFK-down group are related to growth factor binding and receptor signaling (**Figure 4**). Enrichment of these genes suggests that epithelial cells actively contribute to shaping the extracellular matrix and that these tissues differentially organize the extracellular microenvironment. This likely has important consequences for tissue homeostasis.

### An online database allows for easy analysis of differential gene expression

This paper describes a careful transcriptional analysis of 3D organotypic rafts derived from different tissues. To maximize the impact of this resource, it is essential to share these data with the community, both in the form of raw data that allows for reanalysis and in the form of a pre-analyzed database that allows for easy reference. We developed an interactive database using the R statistical language and Shiny app, available at https://viz.datascience.arizona.edu/3DEpiEx/. The website allows users to visualize the expression of individual genes of interest across the different tissues (screenshot **Figure 5A**) as well as in 2D volcano plots (screenshot **Figure 5B**, similar to the plots in **Figure 6**) and an interactive 3D volcano plot linking differential expression and statistical support.

**Figure 5.**
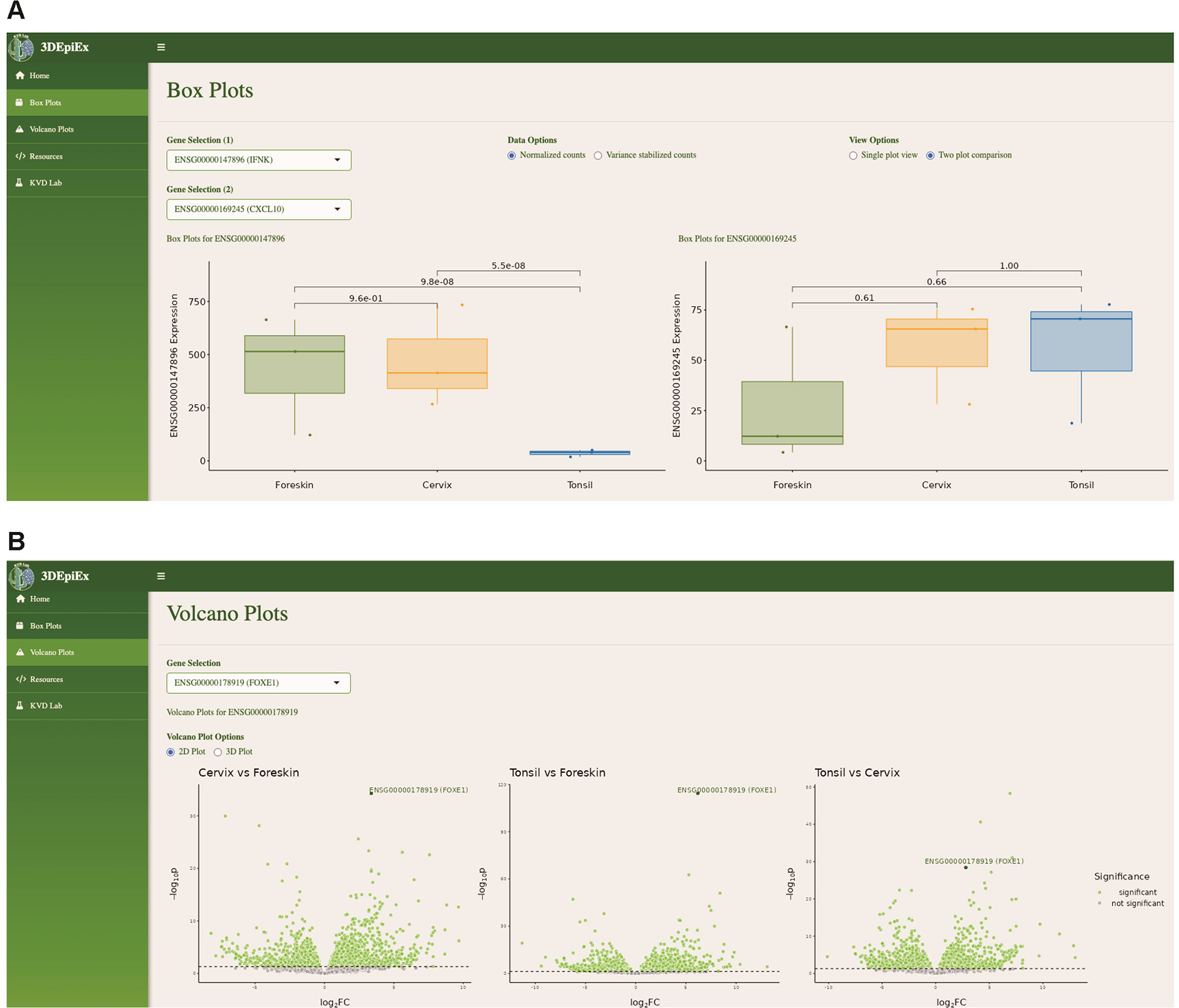
3DEpiEx interactive database. Screenshots of our web-based three-dimensional (<u>3D</u>) <u>Epi</u>thelial Gene <u>Ex</u>pression (3DEpiEx) application which allows users to search genes of interest and visualize their expression in foreskin, cervix, and tonsil-derived 3D epithelial raft cultures as **(A)** boxplots of normalized or variance-stabilized counts and **(B)** volcano plots in the context of pairwise contrasts.

**Figure 6.**
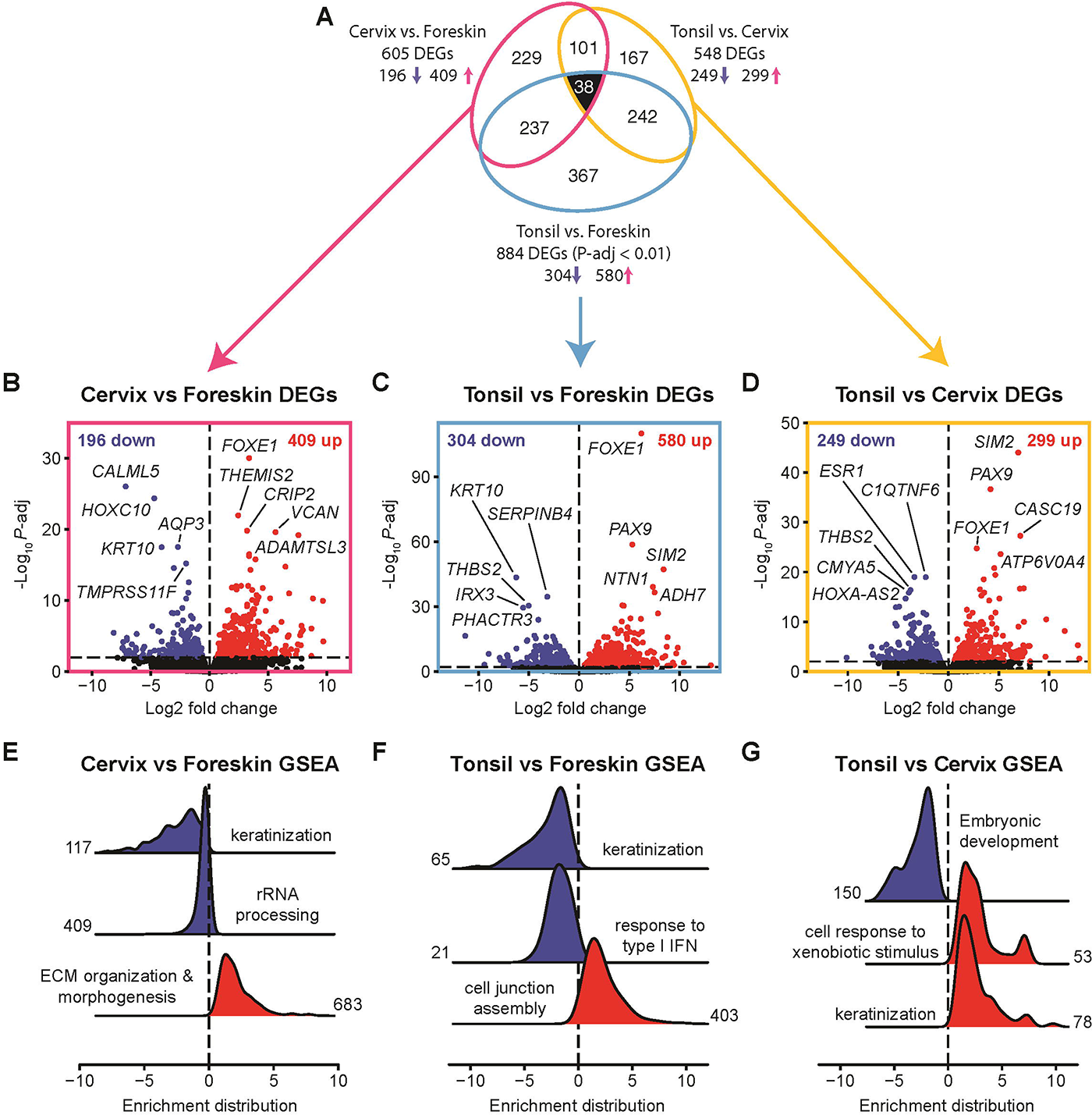
Pairwise contrasts identify the differentially expressed genes between foreskin, cervix, and tonsil-derived 3D epithelia. **(A)** Euler diagram with area-proportional ellipse for each pairwise contrast, including up (red) and down (blue) genes. **(B-D)** Volcano plots of the DESeq2 pairwise contrast results. The top five up-and down-regulated genes are labeled. **(E-G)** Ridge plots of the aggregated core enriched gene distribution (log_2_ fold change) from pairwise gene set enrichment analysis (GSEA): non-redundant functional modules from network analysis of the statistically significant Gene Ontology Biological Process (GO:BP) terms are shown (Figure 6-figure supplements 1-3). The number of core enriched (leading edge) genes are shown beside each distribution. Red = enriched. Blue = depleted.

### Pairwise contrasts between 3D tissues highlight their distinct transcriptional and functional profiles

Due to their availability, primary human foreskin keratinocytes are often used in research (Arnette et al., 2016). Our global analysis identified that the foreskin samples clustered apart from the cervix and tonsil tissues, indicating that these skin-derived cells are phenotypically and transcriptionally distinct from each of the mucosal-origin tissues studied here. Indeed, the two primary clusters of differentially expressed genes corresponded to down- or up-regulation in the foreskin (**Figure 4**).

We updated the described DESeq2 model to make direct pairwise comparisons between each of the three epithelial tissue origins [cervix vs. foreskin (**Supplementary File 8**), tonsil vs. foreskin (**Supplementary File 9**), and tonsil vs. cervix (**Supplementary File 10**)]. Among the 18,925 confidently detected genes, we found a total of 1,381 pairwise differentially expressed genes (Wald test *P*-adj < 0.01). These 1,381 identified genes were plotted using a three-way area-proportional Euler ellipses diagram (**Figure 6A**).

The Euler diagram in **Figure 6A** illustrates that many identified pairwise differentially expressed genes are shared across pairwise tissue comparisons. Each pairwise comparison was visualized using individual volcano plots. These plots show the relationship between the magnitude of differential expression (x-axis) and statistical significance (y-axis) for each differentially expressed gene (**Figure 6B-D**). Functional processes enriched and depleted for each comparison were identified using gene set enrichment analyses (GSEA). Individual genes often belong to multiple GO terms, leading to overlap between distinct GO terms. To deconvolve this redundancy, we visualized the statistically significant GO terms using enrichment maps and gene-concept networks (**Figure 6–figure supplements 1-3**). These networks organize enriched terms with edges connecting overlapping gene sets to identify functional modules. For each network, we identified functional modules and plotted the distribution of fold change values of all core enriched (also known as leading edge) genes contained within each module (**Figure 6E-G**). Therefore, these ridge plots show the number of aggregated core genes in each module and their differential expression. Red curves show that a set of core genes is enriched, while blue distributions indicate depletion.

#### Cervix-derived epithelia overexpress genes involved in ECM remodeling and deplete keratinization compared to foreskin

We first compared foreskin tissue to cervical tissue. This comparison identified 605 differentially expressed genes: 409 up- and 196 down-regulated (**Figure 6B**). The top significantly up-regulated genes in cervix vs. foreskin were transcription factor *FOXE1*, putative transcription factor *CRIP2*, immune regulator *THEMIS2*, extracellular matrix proteoglycan *VCAN*, and metalloprotease *ADAMTSL3*. The top down-regulated genes were the calcium-binder *CALML5*, transcription factor *HOXC10*, water channel *AQP3*, suprabasal differentiation maker *KRT10*, and transmembrane protease *TMPRSS11F*. Gene set enrichment analysis on these 605 differentially expressed genes identified 86 biological processes (72 enriched and 14 depleted, **Supplementary File 11, Figure 6–figure supplement 1**), with the functional modules including depletion of keratinization and rRNA processing and an enrichment of extracellular matrix (ECM) remodeling and morphogenesis.

#### Tonsil-derived epithelia have enriched junction assembly and depleted IFN responses compared to foreskin

Next, we compared the tonsil-derived epithelia to cutaneous foreskin epithelia. This comparison identified 884 differentially expressed genes: 580 up- and 304 down-regulated (**Figure 6C**). The top up-regulated genes in tonsil vs. foreskin were transcription factors (*FOXE1*, *PAX9*, and *SIM2*), the laminin-related *NTN1*, and the metabolic enzyme *ADH7*. The top down-regulated genes were suprabasal differentiation marker *KRT10*, serine protease inhibitor *SERPINB4*, glycoprotein *THBS2*, transcription factor *IRX3*, and the phosphatase and actin regulator *PHACTR3*. Tonsil vs. foreskin GSEA yielded 23 GO:BP terms (17 enriched and 6 depleted, **Supplementary File 12, Figure 6–figure supplement 2**), with the functional modules including enriched junction assembly, as well as depletion of keratinization and response to type I interferon (**Figure 6F**). These observations further support a difference in baseline interferon response between these epithelial tissues we first observed in **Figure 4**.

#### Tonsil-derived epithelia have enriched xenobiotic response and depleted morphogenesis compared to the cervix

While the mucosal-origin tissues, cervix and tonsil, differ individually from the foreskin, we were also interested in determining what distinguishes the mucosal tissues from each other. We found 548 differentially expressed genes in a pairwise comparison of tonsil to cervical epithelia: 299 up- and 249 down-regulated genes in tonsil relative to cervix (**Figure 6D**). The top up-regulated genes in tonsil vs. cervix included transcription factors *SIM2*, *PAX9*, and *FOXE1,* as well as the RNA gene *CASC19* and the enzyme *ATP6V0A4*. The top down-regulated transcripts were estrogen receptor and transcription factor *ESR1*, complement and TNF-related *C1QTNF6*, *THBS2*, *CMYA5*, and lncRNA *HOXA-AS2*. Tonsil vs. cervix GSEA yielded 23 GO:BP terms (10 enriched and 13 depleted, **Supplementary File 13, Figure 6–figure supplement 3**), with the functional modules including enriched cellular response to xenobiotic stimulus (*e.g.*, genes encoding cytochrome P450 monooxygenases) and keratinization as well as depletion of genes involved in embryonic development (**Figure 6G**).

### Tissue-identifying gene expression profiles highlight differences in embryonic origin and interferon responses

To investigate tissue-specific gene expression, we analyzed the pairwise overlapping regions of the Euler diagram (**Figure 6**). For example, **Figure 7A** shows genes differentially expressed in foreskin-derived raft tissues compared to both cervical and tonsillar tissue (*i.e.*, the overlap between the tonsil vs. foreskin and cervix vs. foreskin comparisons, highlighted in green in **Figure 7A**). We also considered differentially expressed genes in all three comparisons (38 genes in the black center of the Euler diagram in **Figure 6A**). This set of genes is plotted using a scatterplot. In these plots, genes in the top right quadrant are up-regulated in foreskin tissue, while genes that fall in the lower left quadrant are down-regulated in foreskin compared to the other tissues. Transcripts in the other two quadrants are differentially expressed in the three-way overlap but are incongruent between comparisons. For example, the expression of *ESR1* is statistically different in all three comparisons. However, ESR1 is up-regulated in foreskin-derived tissues compared to tonsil, while ESR1 is down-regulated when comparing foreskin to cervical tissues (**Figures 7G and 7H**). These incongruent transcripts are excluded from further analyses. Genes in the top right (*i.e.*, up-regulated) and bottom left (*i.e.*, down-regulated) quadrants are further analyzed using a combination of g:Profiler and network analyses to identify interconnected functional modules.

**Figure 7.**
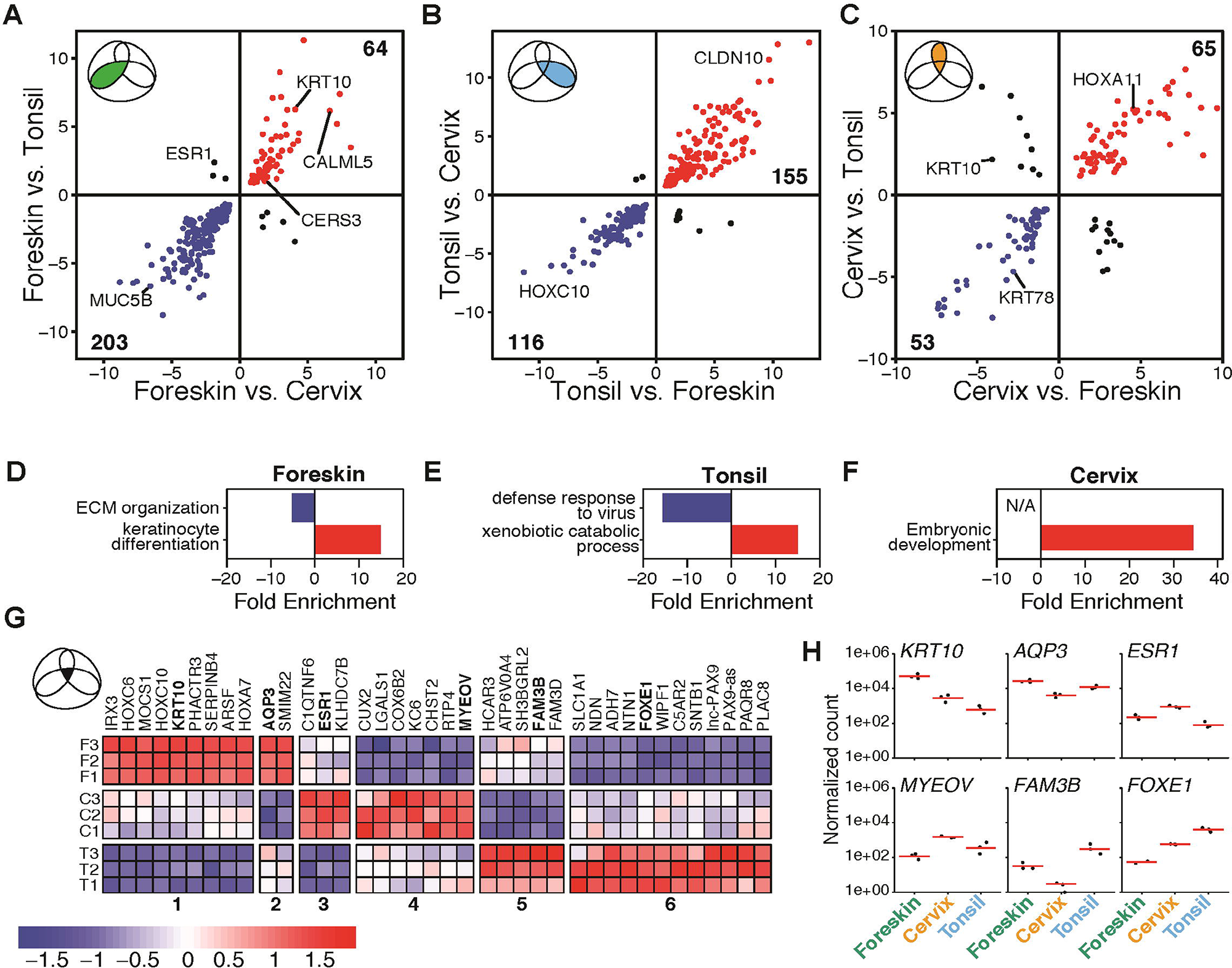
Tissue-identifying differentially expressed genes. **(A-C)** Scatterplots of pairwise contrast overlaps, axes represent log_2_ fold change. **(D-F)** Over-representation analysis enrichment plots for each tissue-identifying gene set, showing the non-redundant functional modules identified via network analysis (**Figure 7–figure supplements 1-3)**. **(G)** Heatmap of the core distinguishing epithelial markers (*n* = 38) of the triple-overlap differentially expressed genes: six groups identified based on tissue expression. The color scale from blue to red represents variance-stabilized, row-centered, counts. **(H)** Log_10_-scaled normalized RNA-seq count data were plotted for genes from each of the six groups.

#### Foreskin-derived rafts uniquely up-regulate genes involved in keratinization

The foreskin-identifying epithelial markers consist of 237 differentially expressed genes plus 30 transcripts found in the Euler diagram’s core representing differentially expressed genes in all pairwise comparisons. Most foreskin-specific differentially expressed genes are down-regulated in foreskin relative to cervix and tonsil (76.0%, 203/267, **Supplementary File 14**). One of the top down-regulated genes in foreskin was *MUC5B*, a gel-forming mucin responsible for lubricating oral and cervical mucosa (Kesimer et al., 2010). Network analysis of enriched functional terms identified a large interconnected functional module (**Figure 7– figure supplement 1**) related to extracellular matrix organization (**Figure 7D**).

Functional enrichment on the 64 up-regulated differentially expressed genes (**Supplementary File 15**) identified 4 terms (**Supplementary File 16, Figure 7–figure supplement 1**) and is defined by keratinocyte differentiation genes (**Figure 7D**). These epidermis development genes include keratinization genes such as regulators of cornification (*e.g.*, *CALML5*, (Méhul et al., 2001)) and ceramide synthases (*e.g.*, *CERS3*). Ceramides are the principal lipids in the stratum corneum and are an essential component of the skin barrier (Murakami et al., 2018).

#### Tonsil-derived rafts uniquely down-regulate baseline antiviral immune responses

When analyzing the oral-derived tonsillar epithelia, we identified 271 (242 + 29) tonsil-identifying differentially expressed genes (**Figure 7B**). 116 of these 271 genes (42.8%) are down-regulated in tonsil-derived tissue (**Supplementary File 17**). Analyzing these genes, we identified 115 enriched functional terms involved in innate immune processes, specifically related to defense responses to viruses and type I interferon signaling, which are down-regulated in tonsillar tissues. These data further support earlier data (**Figure 4**) that the 3D organotypic tonsillar epithelia have a baseline innate immune suppression against viruses (**Figure 7– figure supplement 2,** **Figure 7E**).

In parallel, an analysis of the 155 tonsil up-regulated genes (57.2%, 155/271, **Supplementary File 18**) identified 22 enriched terms (**Supplementary File 19**). Tonsil up-regulated enriched terms related to xenobiotic stimulus and detoxification (*e.g.*, epoxygenase P450 pathway, phenol-containing compound metabolic process, and olefinic compound metabolic process; **Figure 7–figure supplement 2,** **Figure 7E**)

#### Cervix-derived rafts uniquely up-regulate a subset of Hox genes

The 118 cervix-specific epithelial markers include 101 differentially expressed genes from the shared overlap plus 17 core markers (**Figure 7C**). 45% of these differentially expressed genes are down-regulated in cervix compared to the other tissues (44.9%, 53/118, **Supplementary File 20**). The 65 cervix-specific up-regulated differentially expressed genes (55.1%, 65/118, **Supplementary File 21**) were enriched for 115 terms (**Supplementary File 22**) related to embryogenic and morphogenic biological processes (**Figure 7–figure supplement 3**, **Figure 7F**). Of note, several HOX cluster genes are up-regulated in cervical tissue. For example, expression of HOXA11 is critical for the development of the cervix and is strongly up-regulated in these tissues. The down-regulated genes do not appear to be enriched in any functional processes.

### A core set of differentially expressed genes uniquely distinguish each tissue

The three-way overlap identifies 38 differentially expressed genes in all three pairwise comparisons (**Figure 6A**). These 38 genes are differentially expressed in all pairwise comparisons and therefore have a unique expression level for each of the three epithelia. Visualizing the expression of these 38 genes in a heatmap (**Figure 7G**) highlights the different relative expression levels in each tissue. These 38 genes can be grouped into 6 clusters based on relative expression levels. Clusters 1 and 2 contain genes that are up-regulated in tonsils and vary in expression between foreskin and cervix. Clusters 3 and 4 are up-regulated in cervix with differential expression in foreskin and tonsil. Finally, clusters 5 and 6 are up-regulated in foreskin-derived tissues compared to the other 2 tissues.

We highlight 6 genes using plots of the normalized counts to show the magnitude of expression (**Figure 7H**). *KRT10*, a keratin-encoding gene, was among this small subset of genes. *KRT10* was highest in foreskin (Wald *P*-adj = 3.14 x 10^-18^ and 3.17 x 10^-44^, relative to cervix and tonsil, respectively), followed by cervix (Wald *P*-adj = 4.91 x 10^-5^, relative to tonsil), and lowest in tonsil (**Figure 7H**). This keratin is commonly used as an epithelial differentiation marker (Jackson et al., 2020b), found in the spinous layer of cutaneous epithelium (Cheng et al., 2018), but clearly has a tissue-specific abundance.

The stratum corneum of the epidermis is a critical barrier that functions to limit water loss by evaporation. The aquaglyceroporin AQP3 is abundantly expressed in keratinocytes of the mammalian skin epidermis along with the stratified squamous epithelial cells of the oral cavity and esophagus (Agha-Hosseini et al., 2020, p. 3; Sougrat et al., 2002). AQP3 is highest in foreskin compared to both cervix (Wald *P*-adj = 2.9 x 10^-18^) and tonsil (Wald *P*-adj = 1 x 10^-3^), while AQP3 expression is higher in tonsils compared to cervical tissues (Wald *P*-adj = 9.1 x 10^-6^).

The estrogen receptor (*ESR1)* was up-regulated in tissues with a genital origin. Estrogen receptors are highly expressed in the ectocervix (McKinnon et al., 2020) and foreskin (Pask et al., 2008), and we found it was highest in cervix (Wald *P*-adj = 5.82 x 10^-7^ and 9.89 x 10^-20^, relative to foreskin and tonsil, respectively), followed by foreskin (Wald *P*-adj = 1.40 x 10^-3^ relative to tonsil), and lowest in tonsil (**Figure 7H**).

MYEOV (MYEloma Overexpressed gene) was initially identified as a potential oncogene in specific multiple myeloma cell lines (Janssen et al., 2000). Importantly, it has also been implicated in oral squamous cell carcinomas (Janssen et al., 2000). We find that it has the highest expression in cervix (Wald *P*-adj = 3.4 x 10^-12^ and 3.8 x 10^-3^, relative to foreskin and tonsil, respectively), followed by tonsil (Wald *P*-adj = 2 x 10^-3^ relative to foreskin) and foreskin derived tissues.

Family with sequence similarity 3 (FAM3) is a cytokine-like gene family that contains four genes: FAM3A, FAM3B, FAM3C, and FAM3D (Cao et al., 2003). FAM3B (also called PANDER) regulates the epithelial-mesenchymal transition and has been implicated in the development of oral cancer (He et al., 2019). In our data, FAM3B (and its family member FAM3C) are highly expressed in tonsillar cells and lowest in cervix (Tonsil vs. Foreskin: Wald *P*-adj = 1.2 x 10^-7^; Tonsil vs. Cervix Wald *P*-adj = 1.9 x 10^-17^; Foreskin vs. Cervix: Wald *P*-adj = 5.8 x 10^-5^).

Compared to foreskin-derived tissues, *FOXE1* (Forkhead Box E1) is up-regulated in both tonsil (Wald *P*-adj = 8.06 x 10^-111^) and cervix (Wald *P*-adj = 9.04 x 10^-31^). *FOXE1* is a member of the forkhead family of transcription factors. *FOXE1* is primarily considered a thyroid transcription factor that plays a role in thyroid morphogenesis (Fernández et al., 2015). Within the mucosal tissues, *FOXE1* is further up-regulated in tonsils (Wald *P*-adj = 1.69 x 10^-25^) relative to the cervix (**Figure 7H**). During embryogenesis, the Pharyngeal endodermal pouches give rise to tissues responsible for forming the middle ear cavity and eustachian tube, palatine tonsils, thymus, parathyroid glands, and parafollicular cells of the thyroid, potentially explaining the observed expression profile of *FOXE1* in tonsils.

### In vitro-derived tissue-distinguishing gene signatures are shared with in vivo tissues

In the previous sections, we used our 3D organotypic rafts to characterize tissue-identifying gene expression signatures (**Figure 7**). If these *in vitro* tissues (and thus their gene expression signatures) are equivalent to the *ex vivo* samples, we hypothesized that these *in vitro* gene expression signatures should allow for the identifification of the *ex vivo* samples.

The Genotype-Tissue Expression (GTEx) project is a resource database containing tissue-specific RNA-seq data primarily obtained from low-post-mortem-interval autopsies (GTEx Consortium, 2013). We downloaded gene-level read counts of 2,183 samples (skin, oral, or cervical epithelial) from GTEx. Six hundred four samples were reported to be “suprapubic skin” (not sun exposed), 701 samples were “lower leg” skin (sun-exposed), 9 samples were “ectocervix,” 10 samples were “endocervix,” 142 samples were “uterus,” 555 samples were “esophagus mucosa,” and 162 samples were “minor salivary gland” (from the inner surface of the lower lip). For each of these datasets, we used the “tissue-distinguishing” genes plotted in the top right and bottom left quadrants of **Figure 7** to calculate rank-based single-sample signature scores. The higher this score, the more the sample resembles our 3D organotypic tissues (indicated by the dotted line). When using the *in vitro* foreskin-derived signature, 99.2% of the GTEx samples identified as “skin” have positive scores similar to those observed for the *in vitro* tissues (dotted line). On the other hand, 84.3% of the other samples scored negatively. Notably, while some of the other tissues score positive on this scale, they do not compare to the magnitude of the “skin” tissue scores (**Figure 8A**). We observed similar results for the *in vitro* cervix-derived signature, which positively identified 97.5% of uterine, ectocervical, and endocervical samples in our dataset, while 99.9% of the other tissues scored negative (**Figure 8B**). Likewise, our *in vitro* tonsil-derived signature positively identified 98.2% of oral mucosa samples (Esophagus and Salivary gland). Notably, all the other samples scored negative (**Figure 8C**).

**Figure 8.**
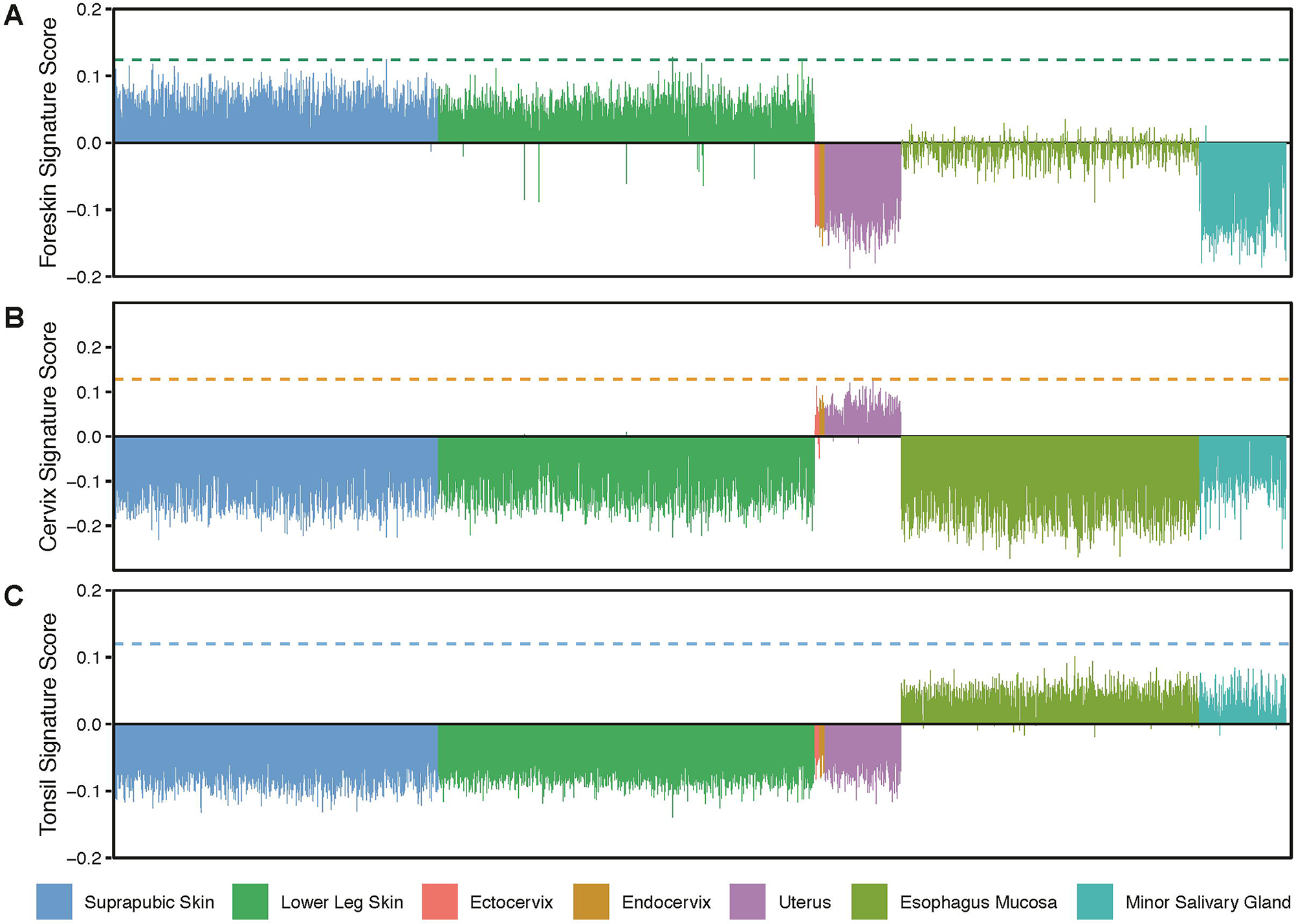
Tissue-distinguishing epithelial signatures identify samples from the GTEx database. Tissue-derived 3D epithelial signature scores were calculated for *n* = 2,183 GTEx RNA-seq samples using our tissue-distinguishing overlap gene sets. We accurately score the samples based on their anatomic site **(A)** foreskin sidentifies skin samples, **(B)** cervix identifies female reproductive tract samples, and **(C)** tonsil identifies oral samples. The dotted line represents the pure epithelial signature score of our 3D raft cultures.

Overall, these findings indicate that our pure 3D epithelial *in vitro* signatures can classify complex *in vivo* tissue samples accurately based on anatomic origin.

### Reduced homeostatic interferon-stimulated gene expression in tonsil cells

Intriguingly, baseline interferon-stimulated gene expression is reduced in tonsillar cells compared to other tissues (**Figure 7E**). We used Integrated Motif Activity Response Analysis (ISMARA) to reconstruct the regulatory networks that control the observed gene expression in these tissues (Balwierz et al., 2014). ISMARA identifies key transcription factors that explain the observed gene expression patterns (Balwierz et al., 2014). The ISMARA analysis identified 4 transcription factor binding motifs with a z-score larger than 2 whose target gene expression does not change between foreskin and cervix but is reduced in tonsil-derived tissue. These transcription factor binding motifs are “IRF2_STAT2_IRF8_IRF1”, “IRF9”, “IRF3”, and “PITX1”. Notably, the baseline expression of these transcription factors is not different between the different tissues (**Figure 9–figure supplement 1**), suggesting that upstream pathways, and not just transcription factor abundance, must be differentially regulated in tonsillar epithelial cells.

A recent study shows an important contribution of STAT2–IRF9 complexes for the constitutive expression of interferon-stimulated genes in resting cells (Platanitis et al., 2019). This subset of interferon-responsive genes is involved in a homeostatic response and is known as ‘tonic interferon-sensitive’ genes. These tonic interferon-sensitive genes were characterized by Mostafavi and colleagues in B-cells and macrophages (Mostafavi et al., 2016). We reanalyzed the Mostafavi dataset and classified their set of interferon-stimulated genes (*n* = 188) as having high (27 genes), medium (15 genes), or low (146 genes) tonic sensitivity (**Figure 9A**). Genes with high tonic sensitivity have a higher baseline of unstimulated expression compared to medium-sensitive genes. For example, IFIT3 and SLFN5 are “high” tonic-sensitive genes; IFIT3 is differentially expressed in tonsillar epithelia, while SLFN5 does not reach statistical significance.

**Figure 9.**
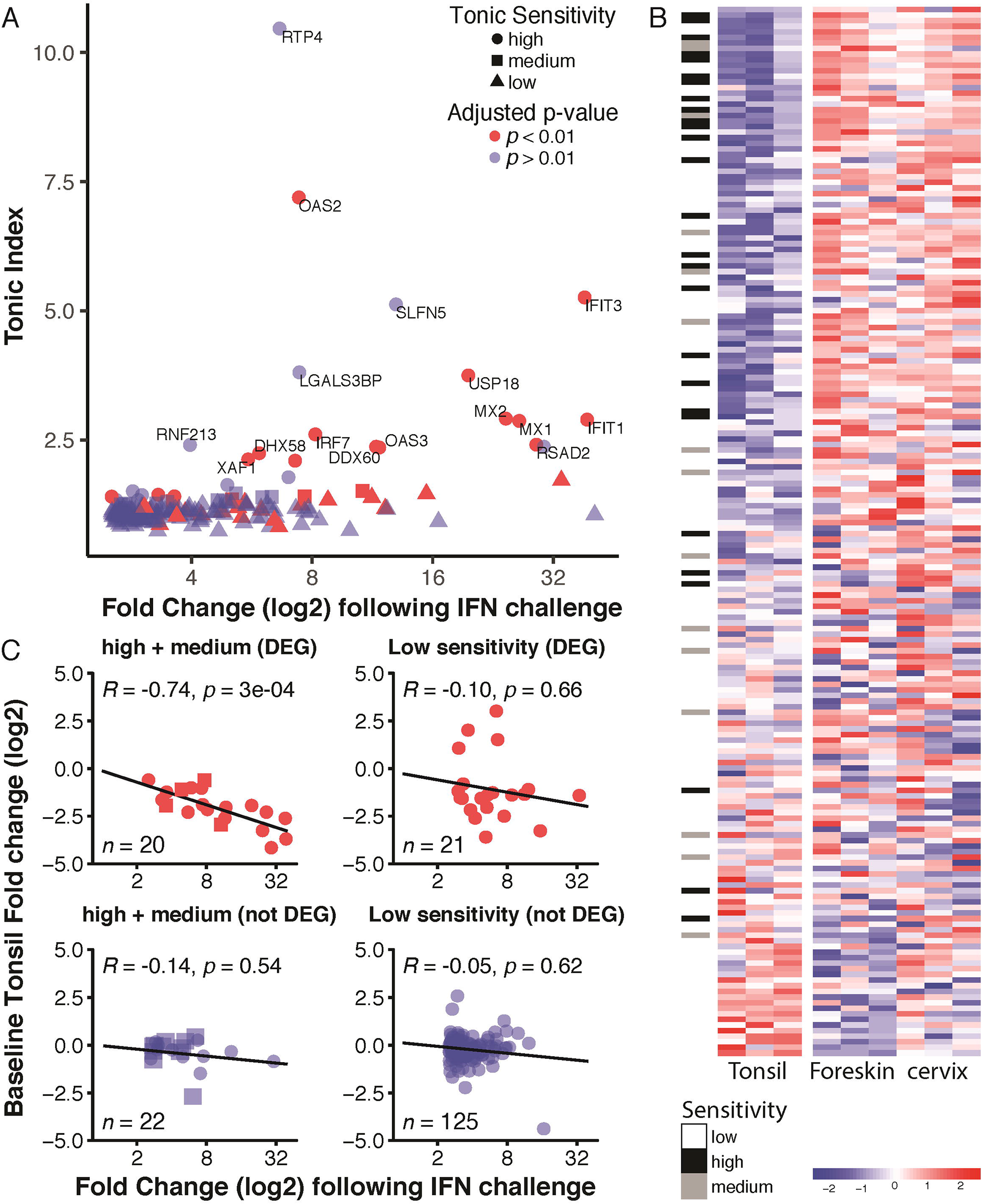
Tonsilar epithelia down-regulate tonic-sensitive interferon genes. **(A)** Scatterplot showing Interferon Stimulated Gene expression vs. dependence on baseline tonic interferon signaling (data aggregated from (Mostafavi et al., 2016)). For details on how data was reanlyzed and aggregated, see materials and methods section. Symbols indicate tonic sensitivity (circle: high, square: medium, and triangle: low), while colors indicate whether the gene is significantly (red: *p*<0.01, blue: *p*>0.01) differentially expressed in tonsils vs. cervix and foreskin. **(B)** Heatmap of the Mostafavi gene set with rows (genes) sorted by fold change of tonsils relative to the other two epithelia. The black and grey bars indicate high and medium tonic sensitivity, respectively. **(C)** Correlation plots of genes with either tonic sensitivity [high (circles) + medium (squares)] or low sensitivity, stratified by whether genes are differentially expressed in tonsil relative to the other two epithelia. R-squared value and p-value for the linear regression line are shown. N indicates the number of genes in each set.

Similarly, 11/27, 4/15 medium, and 21/146 low sensitivity genes are differentially expressed in tonsils. We used a Chi-square test to determine that differentially expressed genes are enriched among high vs. low sensitivity tonic genes (*P* < 0.0001). Likewise, high-sensitivity genes are more likely to be differentially expressed than genes with medium sensitivity (*P* = 0.043). This suggests that tonic-sensitive interferon-responsive genes are specifically down-regulated in tonsils. To further investigate this connection, we plotted the Mostafavi gene set (*n* = 188) using a heatmap. The rows (*i.e.*, genes) in this heatmap are sorted by fold change comparing tonsils to the other two epithelia, with the gene with the biggest reduction (CCL2) on the top and the gene with the largest increase in expression (GBP6) on the bottom. Tonic-sensitive groupings are indicated to the left of the heatmap. The top 10% down-regulated genes (*n* = 18) contain 8 highly sensitive, 2 medium, and 8 lowly sensitive genes. Therefore, the top 10% of down-regulated genes contain 23.8% of the tonic-sensitive (high and medium) genes. Our data suggest tonic-sensitive genes are specifically down-regulated in tonsillar epithelia, potentially limiting interferon responses due to constant exposure to viruses and microbes in the oral cavity (Abt et al., 2012). Following that logic, we hypothesized that tonsillar epithelia might specifically reduce those tonic-sensitive genes that have the highest induction following interferon stimulation. For each gene in **Figure 9B**, we plotted response intensity after interferon treatment (derived from (Mostafavi et al., 2016)) against baseline expression in tonsils (fold change vs. other epithelial tissues). For statistically differentially expressed genes with high (circles) or medium (squares) tonic interferon sensitivity (**Figure 9C**), tonic sensitivity is inversely correlated with baseline expression. This suggests that, in tonsillar epithelia, genes that are highly stimulated by exposure to interferon are more repressed at baseline. No correlations were observed for genes with low tonic sensitivity. Taken together, these data suggest that tonsillar epithelia down-regulate a subset of interferon-sensitive genes, and this repression is correlated with the gene’s expression following interferon stimulation.

## DISCUSSION

For this study, we used a reductionist approach, focusing on the epithelial cells comprising stratified squamous epithelial barriers. While modifications can be made to these 3D models depending on the experimental context and complexity desired, we used an organotypic 3D raft model with the same fibroblasts, collagen, media, growth conditions, and duration to maximize our ability to compare between tissues. For example, 3D organotypic skin and cervical models have been embedded with additional cell types, such as Langerhans cells (Jackson et al., 2020a; Kosten et al., 2015), and others have tested tissue-specific hormone supplementation (estrogen and progesterone) when culturing 3D organotypic ectocervical epithelium (McKinnon et al., 2020). A primary goal of our study was to provide evidence that these different *in vitro* tissues maintain similarities with their *in vivo* tissue equivalents, independent of other cell types or microbiota normally present in these tissues.

Our study used a strict statistical cutoff for identifying differentially expressed genes. Therefore, we may have missed minor (but biologically relevant) differences between the tissues. However, the identified transcripts are likely highly biologically relevant. Aggregating the pairwise contrast data yielded slightly more differentially expressed genes than the global likelihood ratio test: 1381 pairwise vs. 1238 global (see **Supplementary File 23** for the combined global and pairwise results). Comparing the pairwise vs. global differentially expressed genes, 1,194 (83.8%) were pairwise and global differentially expressed. 44 (3.1%) were only global differentially expressed genes, and 187 (13.1%) were only pairwise differentially expressed genes. These differences between the sets of pairwise and global differentially expressed genes are due to the different statistical methods (Love et al., 2014) underlying each test (global likelihood ratio test across all three sample groups vs. pairwise Wald tests between two sample groups) and the high stringency cutoffs (*P*-adj < 0.01) resulting in variable statistical power for detecting differentially expressed genes (Yu et al., 2017).

### Donor-derived epithelial tissues retain origin-specific differences and are valid models

Overall, we found that 3D organotypic epithelia retain their origin-specific characteristics, with anatomic origin sites dominant over donor variability. This was evident at the tissue (histology and immunofluorescence) and transcriptional levels. Overall, many of the tissues’ differences are related to homeotic genes. These genes, *e.g.*, HOX and PAX cluster members, are master regulators directing the development of particular body segments or structures (Gehring, 1993). We found homeotic genes among the top differentially expressed genes in each pairwise contrast, with *HOXC10* highly expressed in foreskin-derived epithelia, *HOXA11* in cervix-derived epithelia, and *PAX9* in tonsil-derived epithelia. *HOXC10* is expressed during mouse urogenital development (Hostikka and Capecchi, 1998) in the analogous tissues that would give rise to the human foreskin. *HOXA11* is expressed in the developing cervix (Taylor et al., 1997). *PAX9* is essential for tissues developing from the pharyngeal pouches (Peters et al., 1998), including the palatine tonsil. These *in vitro* 3D epithelial tissues retain the expression of tissue-specific homeotic regulators, supporting the utility of using epithelial cells from distinct origins in 3D models, implying that these regulatory genes may be valuable biomarkers for distinguishing tissues and verifying their provenance. Importantly, we used gene expression signatures derived from these *in vitro* 3D tissues to classify samples from the GTEx database. Therefore, these pure 3D tissues maintain many transcriptional and histological features of complex *ex vivo* tissue samples.

### Foreskin-derived cells are imperfect models for mucosal tissues

Due to the relative ease of access to foreskin-derived keratinocytes (Choi et al., 2017), these cells have become the *de facto* gold standard of stratified squamous epithelial tissue culture. However, our data clearly demonstrate that, depending on the research question, there may be better models than these cells. Indeed, based on global gene expression analysis, mucosal tissues are transcriptionally distinct from foreskin-derived tissues. This further supports efforts by us and others to transition to more relevant tissues during experimentation (Chatterjee et al., 2019; Doerflinger et al., 2014; Hjelm et al., 2010; Israr et al., 2018; Jackson et al., 2020b; Łaniewski and Herbst-Kralovetz, 2021; Roberts et al., 2019).

Based on gene ontology analyses, many of the differences between tissues are related to “keratinocyte differentiation.” Nonetheless, our histology and immunofluorescence data clearly demonstrate that all three tissues support terminal differentiation.

The Gene Ontology Consortium attempts to create a nomenclature describing how a gene product may regulate biology. Almost by definition, GO annotations rely on the source and data to support an asserted association between a gene product and a GO definition (Balakrishnan et al., 2013). Therefore, since most research on keratinocyte differentiation is based on skin-derived cells, GO terms related to “keratinocyte differentiation” will be biased to skin-derived tissues and will, artefactually, propose that differentiation and associated terms are reduced in these mucosal tissues. Studies like these focused on characterizing these tissues can help update and continue to improve Gene Ontology and related resources.

Additionally, differences within specific epithelia, such as tonsil epithelia, further distinguish epithelia from one another. Clearly, utilizing foreskin-derived cells or 3D epithelia to study host-pathogen interactions or diseases that target specific epithelia elsewhere in the body may not accurately reflect the impact those interactions have on specific genes that are expressed or under-expressed in that particular epithelia.

### Tonsil cells have a reduced baseline immune response

Tonsils are a part of the mucosa-associated lymphoid tissue (MALT), protecting the entrance to the respiratory and alimentary tract (Hellings et al., 2000). As such, tonsils are continuously exposed to microbiota and pathogens. Cells and tissues must balance homeostasis and inflammation. Therefore, tonsils must be able to discriminate between potentially infectious pathogens and innocuous airborne and food antigens. We demonstrate that tonsillar epithelia have reduced baseline expression of interferon-sensitive genes compared to cervical and foreskin tissues. Specifically, this reduction is predicted to be driven through STAT2 and IRF9. Upon type I interferon activation, STAT1, STAT2, and IRF9 form the ISGF3 complex to drive the expression of the full spectrum of interferon-stimulated genes. However, preformed STAT2-IRF9 complexes control the basal expression of interferon-stimulated genes (Platanitis et al., 2019). This basal interferon expression is known as tonic interferon sensitivity (Gough et al., 2012, 2010; Mostafavi et al., 2016). We show that tonsil cells down-regulate these tonic-sensitive genes’ baseline expression. Furthermore, those genes that are strongly induced upon exposure to interferon are the most strongly repressed, presumably increasing the activation energy to lead to transcriptional activation. Of note, STAT2 or IRF9 are not differentially expressed in tonsils, suggesting that other mechanisms must be regulating these pathways. The nature of these control mechanisms remains to be determined but appears to be inherent to explanted tonsil cells as it does not require interaction with other cells or microbes. It is tempting to hypothesize a role for chromatin modifications (Ni et al., 2005).

These data suggest that tonsillar epithelia may have a reduced sensitivity to interferon. Interestingly, it was demonstrated that the microbiota continuously induces low-level interferon and that this baseline stimulation partially regulates the tonic activity of the interferon signaling pathways (Abt et al., 2012). Since tonsils are constantly exposed to microorganisms in the oral cavity, it is tempting to speculate that reduced sensitivity to type I interferon could be beneficial. Interestingly, high concentrations of type I interferons may block B-cell responses (McNab et al., 2015). During an adaptive immune response, germinal centers develop within the tonsils. B cells congregate in the germinal centers and differentiate into long-lived antibody-secreting plasma cells and memory B cells. While interferon stimulates the development of germinal centers, dysregulated germinal center responses may lead to the development of auto-immune disease (Baechler et al., 2003; Bennett et al., 2003; Kirou et al., 2005; Pollard et al., 2013). Indeed, interferon signaling and improper B-cell activation may promote lupus pathogenesis (Jackson et al., 2016). Furthermore, it is well known that excessive interferon signaling has immunosuppressive effects and/or causes inflammation and tissue damage that exacerbate disease. Our data suggest that tonsil epithelia evolved to shape the local immune environment to avoid excessive immune activation.

Indeed, mice without a microbiome due to antibiotic treatment do not mount appropriate innate and adaptive antiviral immune responses following exposure to different viruses (Abt et al., 2012; Invernizzi et al., 2020). Furthermore, these mice had severe bronchiole epithelial degeneration after influenza virus infection (Abt et al., 2012; Lee et al., 2019), implicating an important role for tonic interferon signaling in balancing inflammation and immune responses. Indeed, it was previously demonstrated that tonsillar dendritic cells demonstrate a reduced type I interferon response to influenza infection compared to dendritic cells isolated from blood (Vangeti et al., 2019). Remarkably, different bacteria (*S. aureus, H. influenzae, and N. meningitidis*) activate a robust type I interferon response in freshly purified blood dendritic cells (Michea et al., 2013). However, when the same dendritic cells were exposed to bacteria in media conditioned by tonsillar epithelia, the interferon response decreased 6-fold compared to control media (Michea et al., 2013). This suggests that tonsillar epithelia produce paracrine (and potentially autocrine) factors that regulate the interferon response in dendritic cells. Importantly, these tonsillar epithelial “educated” dendritic cells still support T-cell differentiation and T-cell-derived cytokine production (Michea et al., 2013). This suggests that tonsillar epithelia signal dendritic cells to become less inflammatory in response to constant microbiota exposure while supporting robust adaptive immunity. However, unlike what was proposed by the authors (Michea et al., 2013; Wu et al., 2011), we do not think that reprogramming the local immune microenvironment is a shared feature of all mucosal epithelia. Our data suggest that tonsillar epithelia may be unique in this by reducing the baseline expression of specific interferon-stimulated genes.

This differential regulation of interferon-stimulated genes may partly explain a puzzling result. Chatterjee and colleagues used microarrays to compare the gene expression profile of different tissues infected with HPV16 (Chatterjee et al., 2019). The authors noted that unlike in foreskin-derived tissue, HPV16 did not reduce the expression of immune-related genes in tonsil-derived tissue. However, this study was not designed to directly compare uninfected foreskin and uninfected tonsil tissue (Chatterjee et al., 2019). Our data suggest that the expression of these immune-related genes is already reduced in tonsils. Importantly, this raises the possibility that papillomaviruses (and other pathogens) may differentially interact with tonsils vs. other tissues. This is especially interesting in light of the increasing incidence of papillomavirus induced oropharyngeal (*i.e.,* tonsil and base of tongue) cancer (Chaturvedi et al., 2011).

### Origin-specific epithelial models to study host-pathogen interactions

These differences in tissue-specific gene expression may be important to consider when studying host-pathogen interactions, as they may influence the susceptibility and response of these tissues to infection. Epithelial tissues are a niche for pathogenic microbes, as well as commensals, including a broad range of epitheliotropic fungi, parasites, bacteria, and viruses. Many of these microorganisms have a very specific tropism for particular epithelia. For example, the dermatophyte fungi that cause ringworm thrive in the terminally differentiated apical layers of cornified epithelium and do not colonize host mucosa (Weitzman and Summerbell, 1995). In contrast, other microorganisms, such as specific human papillomaviruses, exhibit tropism toward mucosal rather than cutaneous sites (and vise versa). Importantly, microbes know the difference between specific mucosal stratified epithelia. For example, *Neisseria gonorrhoeae* specifically infects the cervical epithelia, while a related pathogen (*Neisseria meningitidis*), is specific for the oral cavity (Quillin and Seifert, 2018). Similarly, herpes simplex virus type 1 and type 2, primarily infect the genital and oral mucosa, respectively (Garland and Steben, 2014; Löwhagen et al., 2002). Specific papillomavirus types may be even more fastidious, and evolved to infect specific niches on a host epithelia (Egawa et al., 2015; Van Doorslaer et al., 2018). What causes the exquisite tissue tropism seen in these infections is not always apparent, but these specializations do point to biologically critical differences between tissues. The data presented in this resource will allow researchers to generate testable hypotheses explaining these differences in tissue tropism.

## CONCLUSIONS

We characterized biological differences between foreskin, cervical, and tonsillar-derived 3D organotypic epithelial raft cultures using histological, immunofluorescent, and RNA-seq techniques. We confirmed that the identical 3D raft culturing technique yields distinct, origin-specific, stratified squamous epithelia. Epithelial origin is the primary differentiator between these differentiated cultures, yielding distinct transcriptional profiles in the form of thousands of differentially expressed genes. Interestingly, we demonstrate that the transcriptional profile of organotypic tonsillar epithelia is distinguished from foreskin and cervix by a down-regulated baseline expression of important (anti-viral) innate immune genes. Overall, these data are an important resource for those using 3D epithelial models in their research, epithelial biologists, and those studying host-pathogen relationships in these distinct epithelial sites.

## DATA AVAILABILITY

All data is available through the extensive use of supplementary files associated with this manuscript.

The raw data is also available from SRA (BioProject ID PRJNA916745).

## ACKNOWLEDGEMENTS

This work was supported by a National Institute of Dental and Craniofacial Research grant 1R03DE030211-01 and State of Arizona Improving Health TRIF to KVD. We acknowledge fellowship support from the BIO5 Institute and Natural Sciences and Engineering Research Council of Canada (NSERC): Postdoctoral Fellowship (PDF-546182-2020) to RJ. We are thankful to Dr. Aloysius Klingelhutz for primary cervical keratinocytes (donor cultures: C415, C4302, and CX2399), Dr. Deepta Bhattacharya and Dr. Lucas D’Souza for palatine tonsils, and Dr. Felicia Goodrum for HFF-hTERT cells. Thank you to KVD lab members for helpful suggestions, especially David Williams, Isabelle Tobey, Kelly King, and Amy Banka. We also thank Dr. Magdalene So, Dr. Melissa Herbst-Kralovetz, and Dr. Samuel Campos for insightful discussions and critique. Histology services were provided by the University of Arizona Cancer Center’s Tissue Acquisition and Cellular/Molecular Analysis (TACMASR) shared resource core which is supported by the National Cancer Institute of the National Institutes of Health under award number P30 CA023074. The CyVerse Discovery Environment (www.cyverse.org) is a computational infrastructure supported by the National Science Foundation under award numbers DBI-0735191, DBI-1265383, and DBI-1743442.

## SUPPLEMENTARY FILES

**Supplementary File 1.** “Raft_bio_counts.txt”. Tab-delimited gene-level read counts, based on the human reference genome GRCh38, for all RNA-seq samples (*n* = 9).

**Supplementary File 2.** “Raft_bio_groups.csv”. Comma-separated RNA-seq sample metadata including experimental design groups.

**Supplementary File 3.** “Raft_bio_norm_counts.csv”. Comma-separated DESeq2 normalized read counts (library-size corrected).

**Supplementary File 4.** “Raft_bio_global_DEGs_thresh.csv”. Comma-separated DESeq2 likelihood-ratio test results, ordered by lowest *P*-adj.

**Supplementary File 5.** “Raft_bio_global_cluster_annotations.csv”. Comma-separated gene list, ordered by lowest *P*-adj.

**Supplementary File 6.** “Raft_bio_global_HFK-up_ORA.csv”. Comma-separated g:Profiler output, ordered query.

**Supplementary File 7. “**Raft_bio_global_HFK-down_ORA.csv”. Comma-separated g:Profiler output, ordered query.

**Supplementary File 8.** “Raft_bio_pairwise_DEGs_cervix_vs_foreskin_thresh.csv”. Comma-separated DESeq2 Wald test results, ordered by lowest *P*-adj.

**Supplementary File 9.** “Raft_bio_pairwise_DEGs_tonsil_vs_foreskin_thresh.csv”. Comma-separated DESeq2 Wald test results, ordered by lowest *P*-adj.

**Supplementary File 10.** “Raft_bio_pairwise_DEGs_tonsil_vs_cervix_thresh.csv”. Comma-separated DESeq2 Wald test results, ordered by lowest *P*-adj.

**Supplementary File 11.** “Raft_bio_pairwise_GSEA_GO_BP_cervix_vs_foreskin.csv”. Comma-separated ClusterProfiler gene set enrichment results for Gene Ontology Biological Processes. Results ordered by lowest *P*-adj.

**Supplementary File 12.** “Raft_bio_pairwise_GSEA_GO_BP_tonsil_vs_foreskin.csv”. Comma-separated ClusterProfiler gene set enrichment results for Gene Ontology Biological Processes. Results ordered by lowest *P*-adj.

**Supplementary File 13.** “Raft_bio_pairwise_GSEA_GO_BP_tonsil_vs_cervix.csv”. Comma-separated ClusterProfiler gene set enrichment results for Gene Ontology Biological Processes. Results ordered by lowest *P*-adj.

**Supplementary File 14.** “Raft_bio_foreskin_down_genes_203.csv”. Comma-separated gene list of foreskin-identifying down-regulated genes.

**Supplementary File 15.** “Raft_bio_foreskin_up_genes_64.csv”. Comma-separated gene list of foreskin-identifying up-regulated genes.

**Supplementary File 16.** “Raft_bio_overlap_foreskin_ORA.csv”. Comma-separated ClusterProfiler results.

**Supplementary File 17.** “Raft_bio_cervix_down_genes_53.csv”. Comma-separated gene list of tonsil-identifying down-regulated genes.

**Supplementary File 18.** “Raft_bio_cervix_up_genes_65.csv”. Comma-separated gene list of tonsil-identifying up-regulated genes.

**Supplementary File 19.** “Raft_bio_overlap_tonsil_ORA.csv”. Comma-separated ClusterProfiler results.

**Supplementary File 20.** “Raft_bio_tonsil_down_genes_116.csv”. Comma-separated gene list of cervix-identifying down-regulated genes.

**Supplementary File 21.** “Raft_bio_tonsil_up_genes_155.csv”. Comma-separated gene list of cervix-identifying up-regulated genes.

**Supplementary File 22.** “Raft_bio_overlap_cervix_ORA.csv”. Comma-separated ClusterProfiler results.

**SupplementaryFile 23.** “Raft_bio_aggregate_DEGs_thresh.csv”. Comma-separated DESeq2 results including Wald test nominal *P*-values, adjusted *P*-values, and log_2_ fold changes for each pairwise contrast as well as the global likelihood-ratio test nominal *P*-values and adjusted *P*-values for each gene. Results ordered by lowest global adjusted *P*-value.

## FIGURE LEGENDS

**Figure 1–figure supplement 1.**
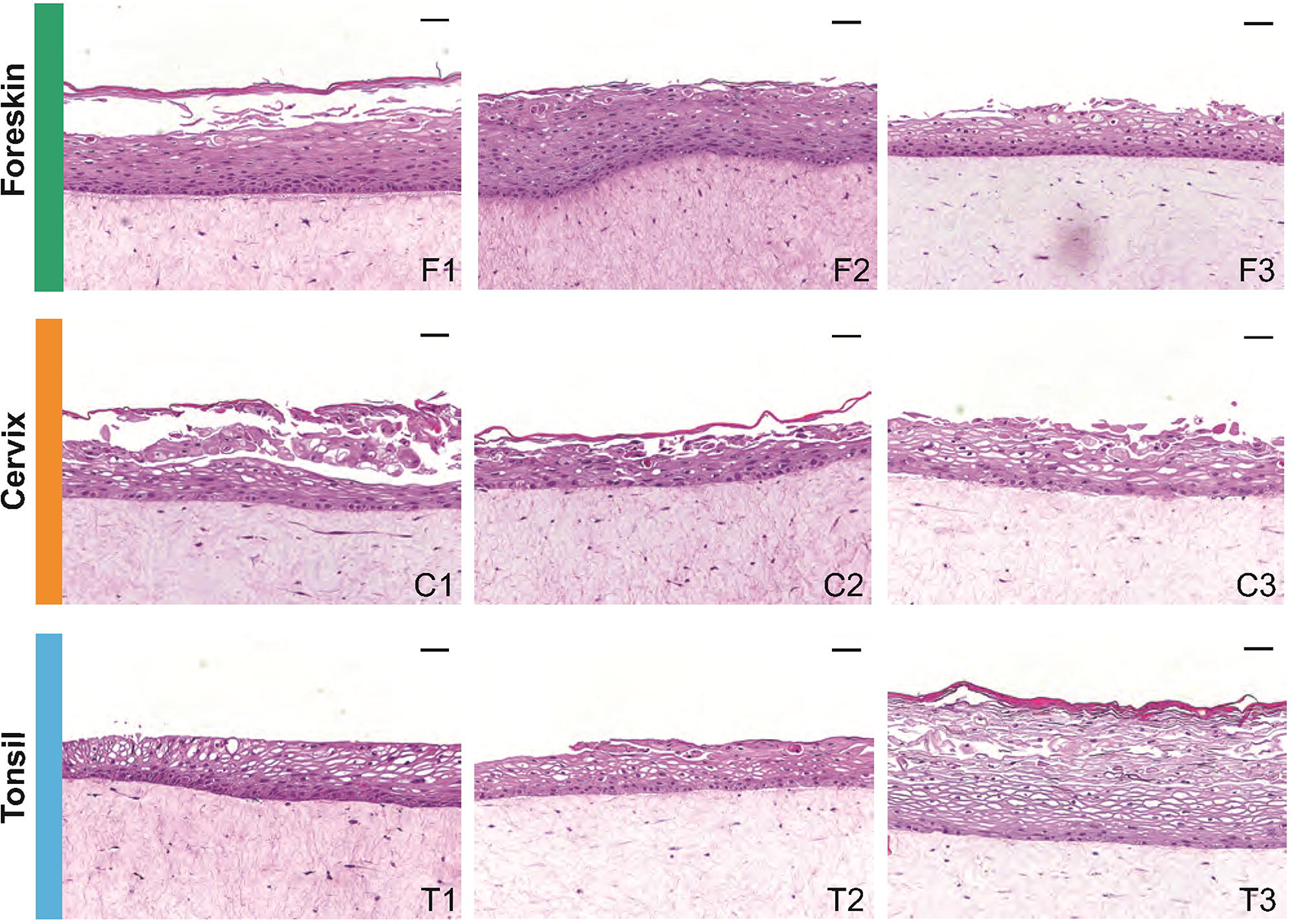
H&E micrographs of all nine independent donor 3D organotypic epithelial raft cultures (*n* = 3 for each tissue origin). Scale bars represent 50 μm.

**Figure 3–figure supplement 1.**
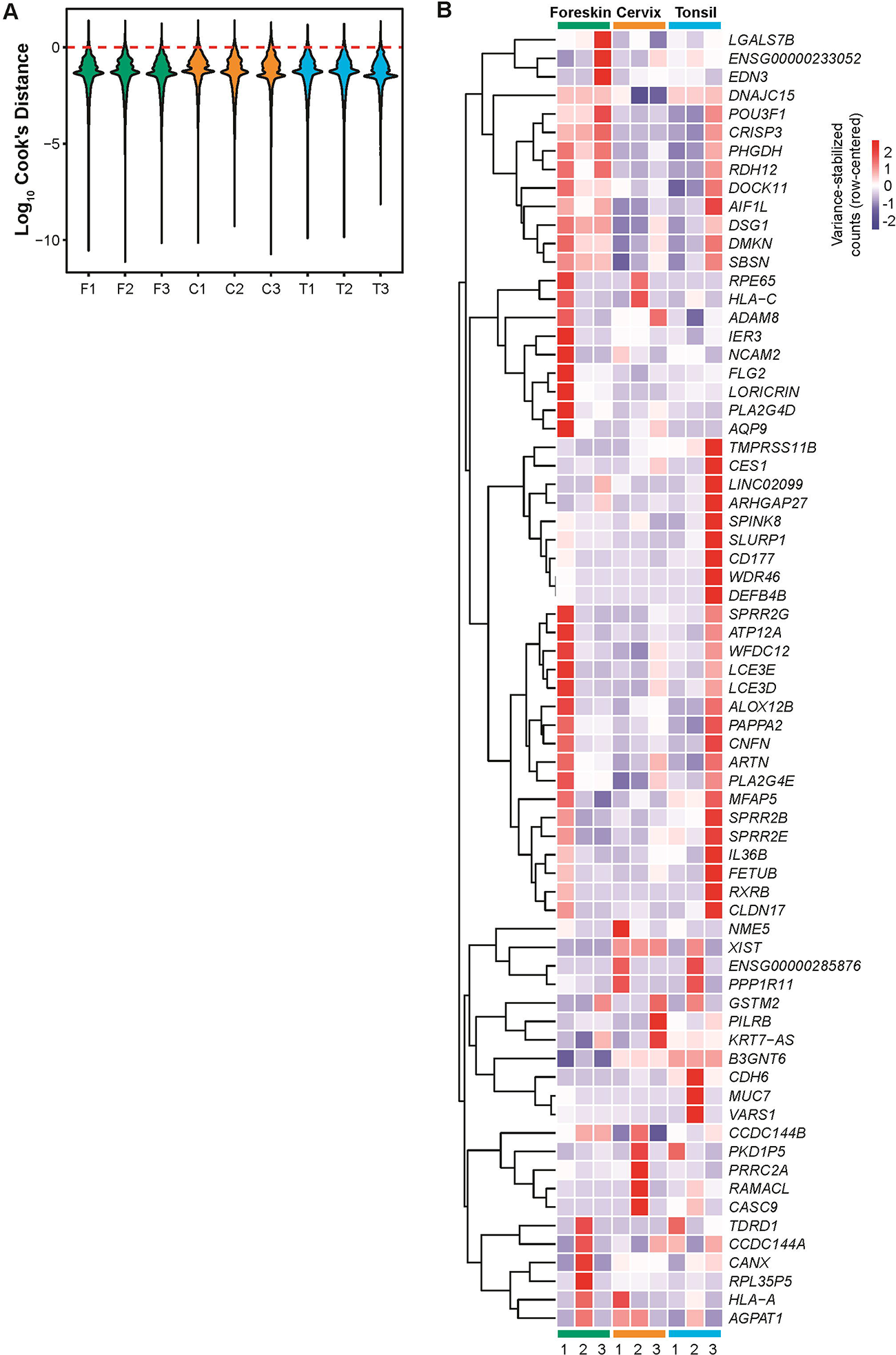
**(A)** Sample outlier quality control: violin plot of log_10_ Cook’s distances (y-axis) to assess if any individual samples (x-axis) were overtly influencing the DESeq2 analyses. All samples were approximately equal and we identified no sample outliers (*i.e.*, violins were all below the red dashed line). **(B)** Heatmap of the 70 outlier genes identified by DESeq2. Scale bar represents row-centered variance-stabilized log_2_ fold change.

**Figure 6–figure supplement 1.**
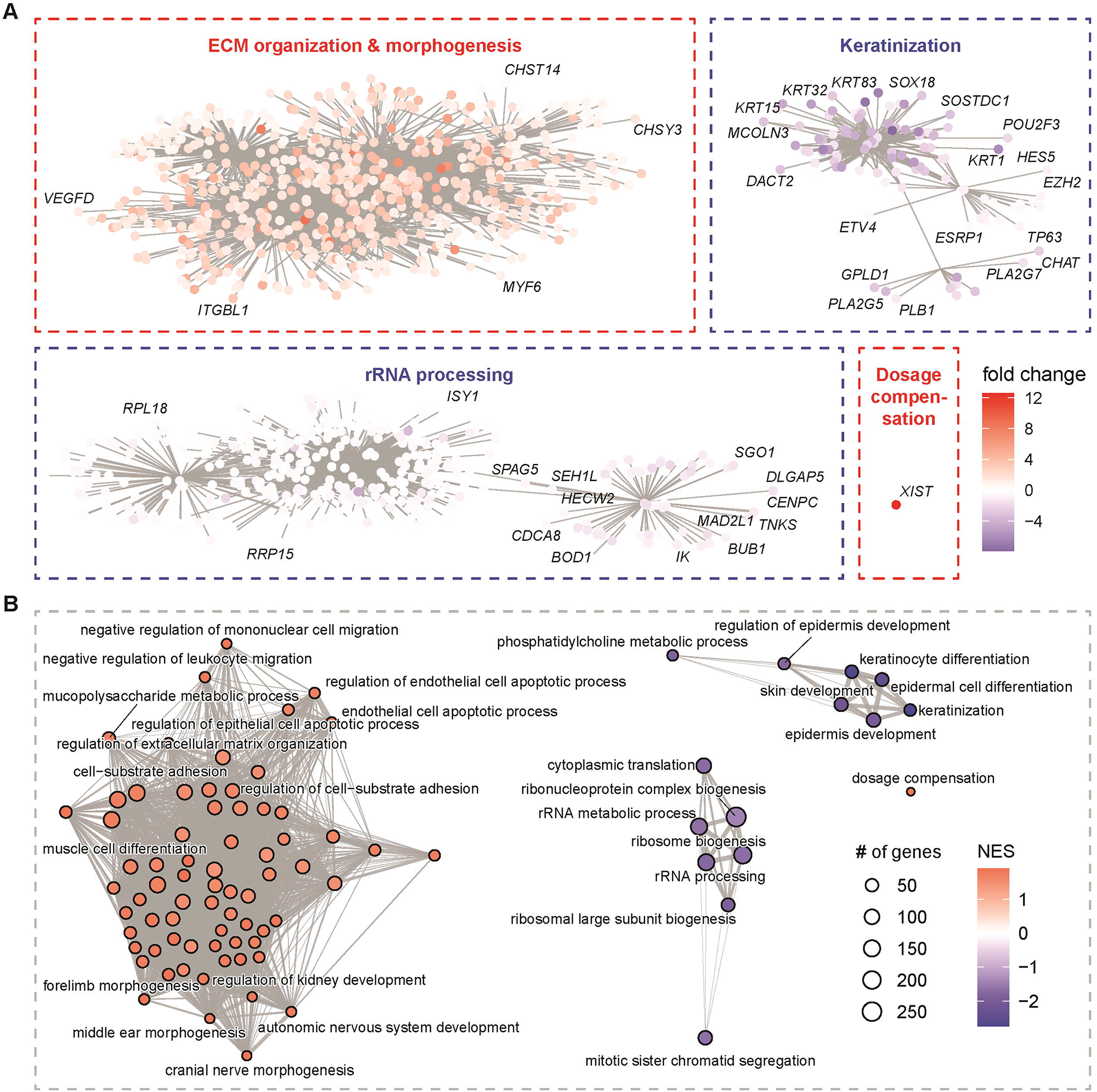
Cervix vs. foreskin functional networks. **(A)** Gene-concept network of the statistically significant Gene Ontology Biological Process (GO:BP) terms from Gene Set Enrichment Analysis (GSEA) of the pairwise contrast results. Nodes are the core enriched genes colored based on their log_2_ fold change (red = enriched and blue = depleted). Connected terms represent functional modules. **(B)** Enrichment map network of just the GO:BP terms colored based on normalized enrichment score (NES).

**Figure 6–figure supplement 2.**
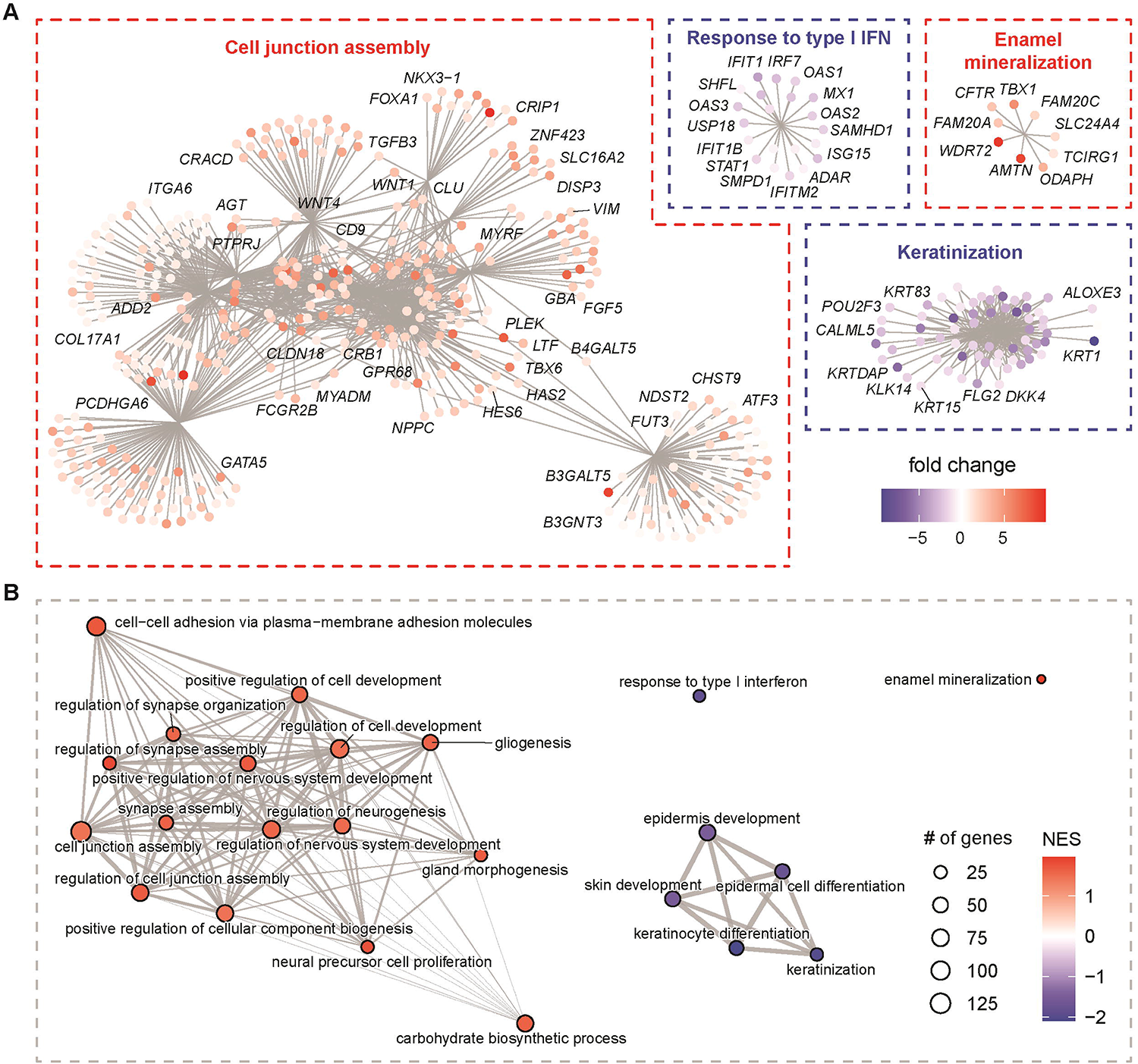
Tonsil vs.foreskin functional networks. **(A)** Gene-concept network of the statistically significant Gene Ontology Biological Process (GO:BP) terms from Gene Set Enrichment Analysis (GSEA) of the pairwise contrast results. Nodes are the core enriched genes colored based on their log_2_ fold change (red = enriched and blue = depleted). Connected terms represent functional modules. **(B)** Enrichment map network of just the GO:BP terms colored based on normalized enrichment score (NES).

**Figure 6–figure supplement 3.**
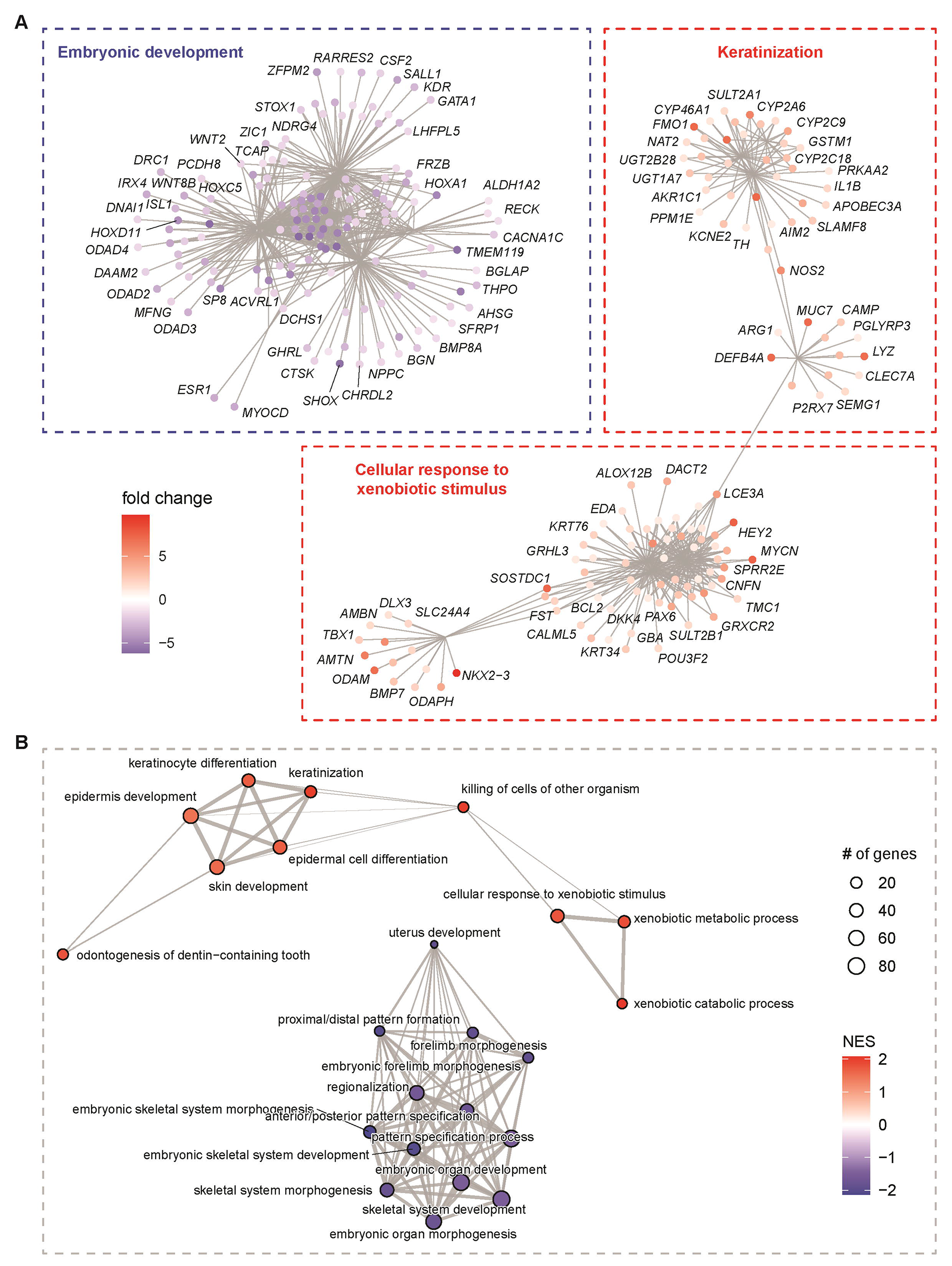
Tonsil vs.cervix functional networks. **(A)** Gene-concept network of the statistically significant Gene Ontology Biological Process (GO:BP) terms from Gene Set Enrichment Analysis (GSEA) of the pairwise contrast results. Nodes are the core enriched genes colored based on their log_2_ fold change (red = enriched and blue = depleted). Connected terms represent functional modules. **(B)** Enrichment map network of just the GO:BP terms colored based on normalized enrichment score (NES).

**Figure 7–figure supplement 1.**
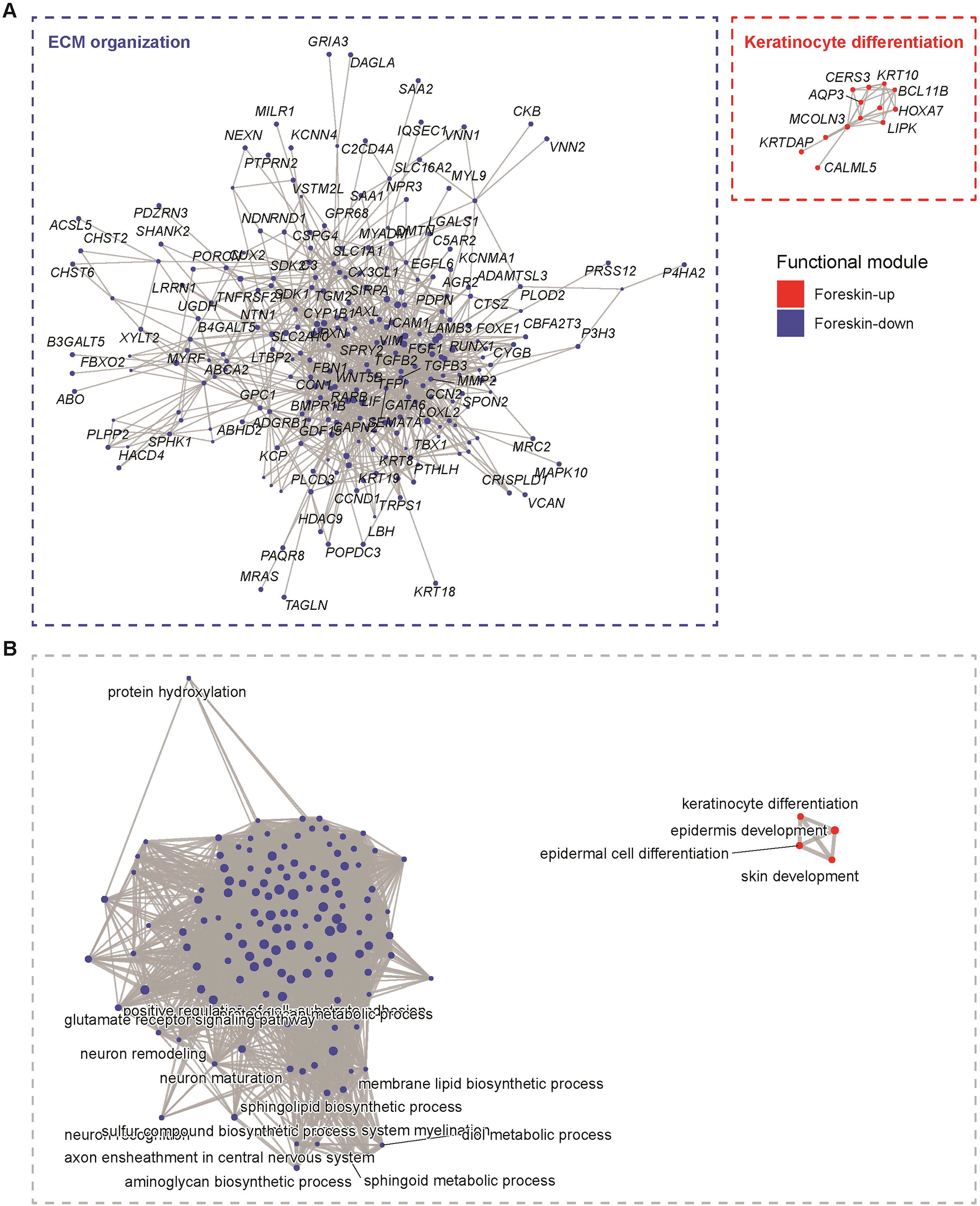
Foreskin-specific functional networks. **(A)** Gene-concept network of the statistically significant Gene Ontology Biological Process (GO:BP) terms from over-representation analysis (ORA) of the tissue-specific differentially expressed genes. Term categories are the hub nodes, while the surrounding nodes are the core enriched genes. Colors: red = up, and blue = down). Connected terms represent functional modules. **(B)** Enrichment map network of just the GO:BP terms colored based on whether they are enriched (red) or depleted (blue).

**Figure 7–figure supplement 2.**
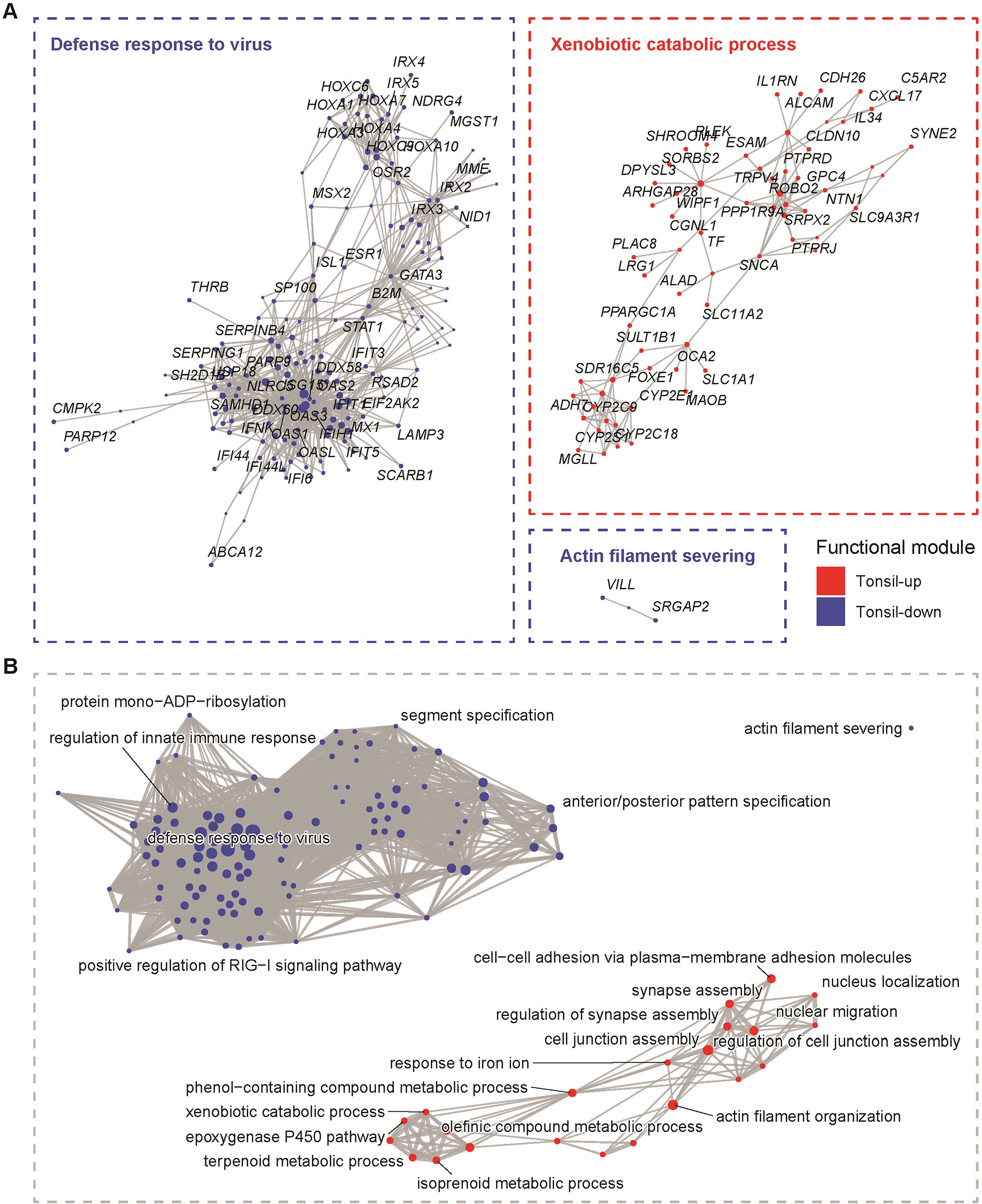
Tonsil-specific functional networks. **(A)** Gene-concept network of the statistically significant Gene Ontology Biological Process (GO:BP) terms from over-representation analysis (ORA) of the tissue-specific differentially expressed genes. Term categories are the hub nodes, while the surrounding nodes are the core enriched genes. Colors: red = up, and blue = down). Connected terms represent functional modules. **(B)** Enrichment map network of just the GO:BP terms colored based on whether they are enriched (red) or depleted (blue).

**Figure 7–figure supplement 3.**
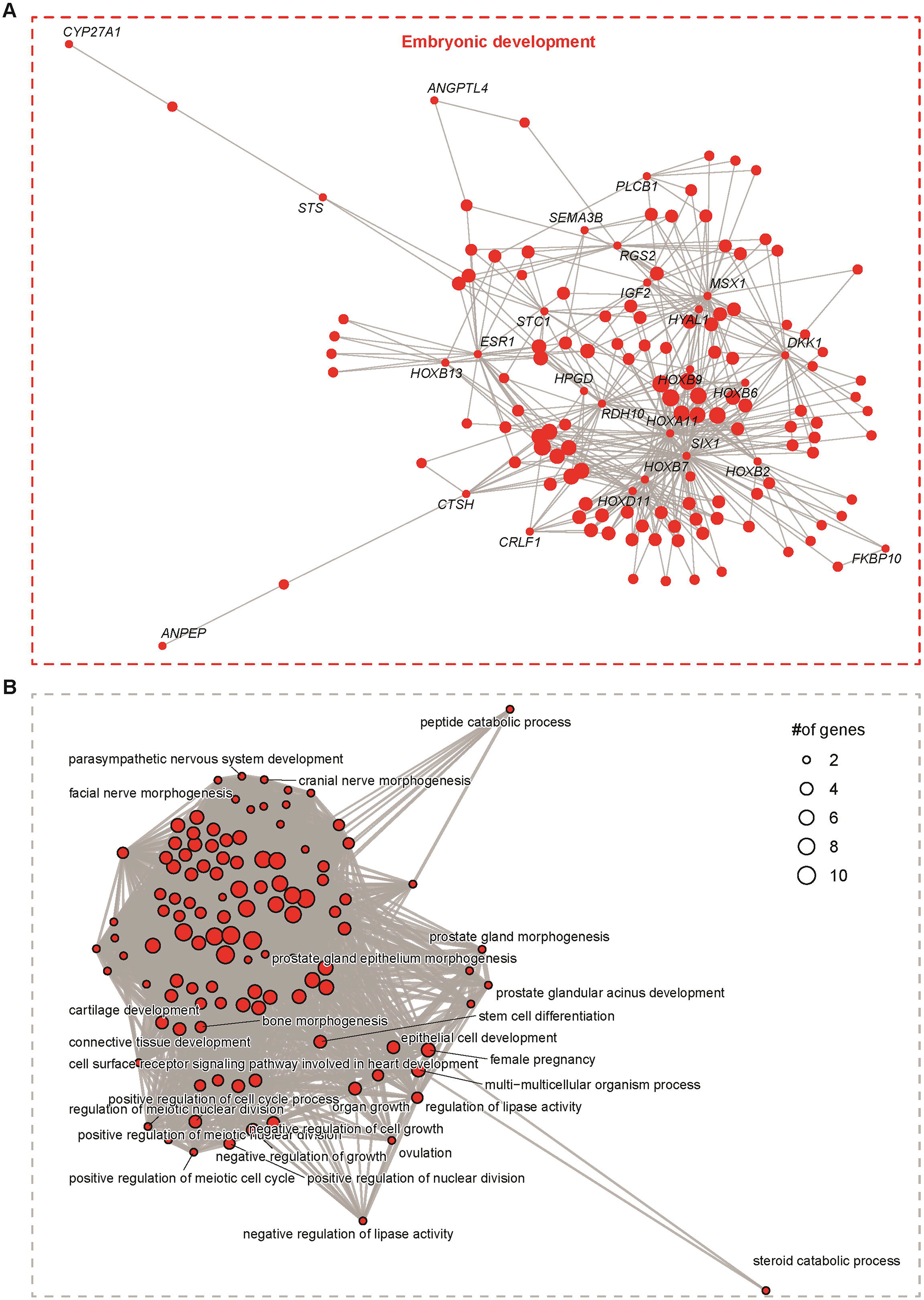
Cervix-specific functional networks. **(A)** Gene-concept network of the statistically significant Gene Ontology Biological Process (GO:BP) terms from over-representation analysis (ORA) of the tissue-specific differentially expressed genes. Term categories are the hub nodes, while the surrounding nodes are the core enriched genes. Colors: red = up). Connected terms represent functional modules. **(B)** Enrichment map network of just the GO:BP terms colored based on whether they are enriched (red).

**Figure 9–supplement 1.**
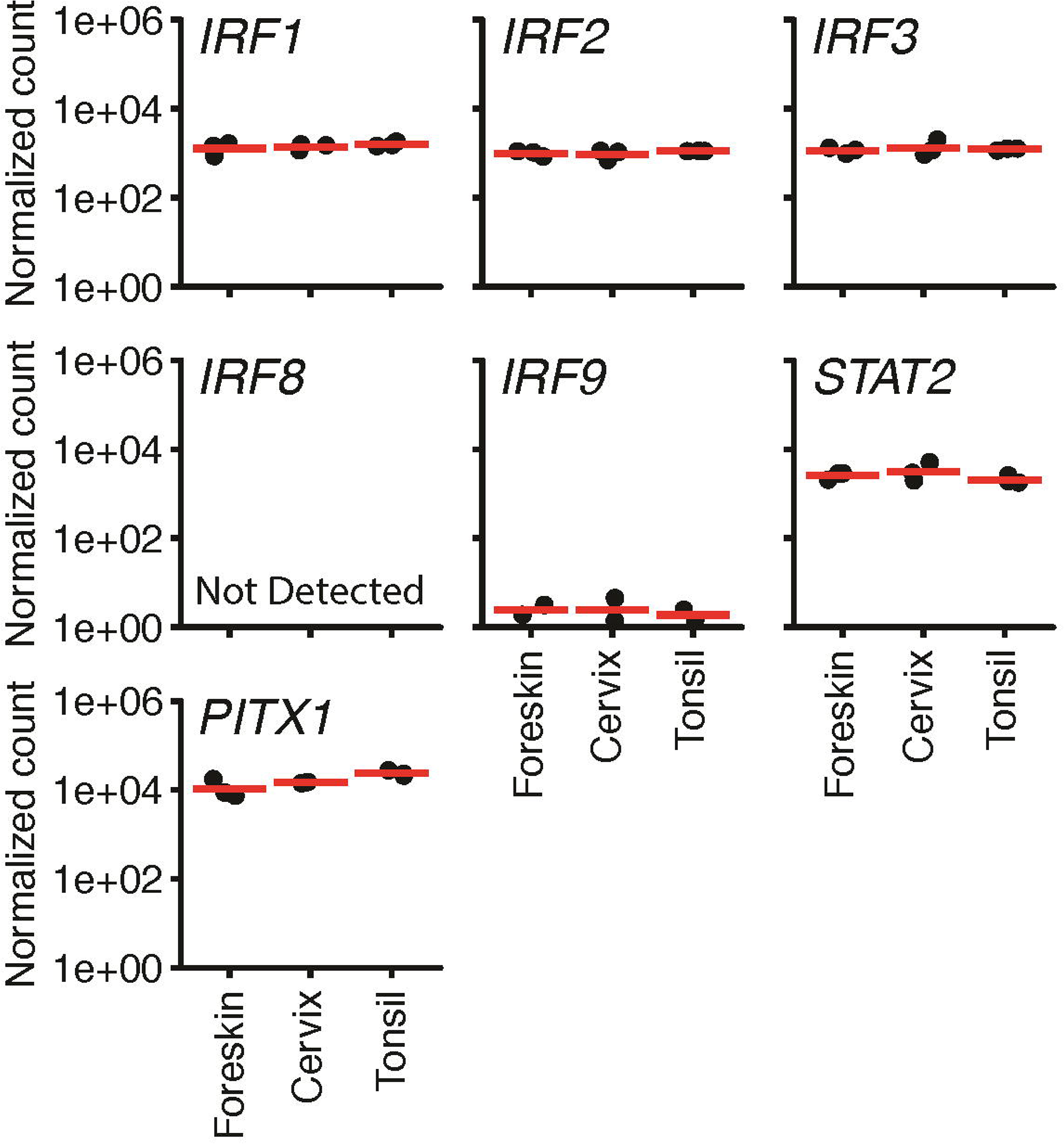
Transcription factors with reduced motif activity in tonsils have unchanged RNA-level expression. Log_10_-scaled normalized RNA-seq count data were plotted for genes encoding transcription factors (and complex components) identified via Integrated Motif Activity Response Analysis (ISMARA): IRF2_STAT2_IRF8_IRF1, IRF9, IRF3, and PITX1. These were binding motifs with a z-score larger than 2 whose target gene expression does not change between foreskin and cervix but are reduced in tonsil-derived tissue.

## Notes

https://viz.datascience.arizona.edu/3DEpiEx/

### Competing Interest Statement

The authors have declared no competing interest.

